# Thalamic stereo-EEG detects sleep stages N2 and N3 with discrimination approaching scalp EEG

**DOI:** 10.64898/2026.10.31.763362

**Authors:** Mahmoud Alipour, Wim van Drongelen, Peter C. Warnke, Naoum P. Issa, Kenneth Lee, David Satzer

## Abstract

Sleep and epilepsy interact strongly during sleep stages N2 and N3, when synchronization of brain rhythms is increased. Sleep staging relies on scalp EEG, which is often absent or incomplete during stereo-EEG monitoring and unavailable from implanted devices, while thalamic leads are increasingly implanted for epilepsy neuromodulation. Given the role of the thalamus in spindle generation and slow oscillation dynamics, we hypothesized that thalamic channels would identify N2 and N3 (N2+N3) better than other intracranial sites. We analyzed 477 h of stereo-EEG with simultaneous scalp EEG from six adults, including 46 thalamic bipolar channels on 11 electrodes. Two classifiers were evaluated on identical channels: a random forest with hidden Markov decoding (RF + HMM) trained leave-one-participant-out, and the pretrained SleepSEEG algorithm developed without thalamic channels. The primary endpoint was discrimination of N2+N3 from other stages as classified from scalp EEG by a sleep expert. Among regions sampled in all participants, thalamic channels performed best (area under the curve [AUC] 0.94 with RF + HMM and 0.93 with SleepSEEG across whole recordings; 0.88 and 0.86 at night). With RF + HMM, thalamic channels outperformed mesial temporal and temporal neocortical channels in all six participants across whole recordings. The advantage persisted against channel-count-matched subsets and a white-matter control with both classifiers. Individual thalamic electrodes reached AUCs of 0.79-0.99 with RF + HMM (10 of 11 at least 0.89). These offline findings support thalamic N2+N3 detection and motivate real-time evaluation for sleep-targeted neuromodulation.

## 1 Introduction

Sleep and epilepsy interact in both directions. Interictal epileptiform discharges and seizures are often facilitated during NREM sleep, particularly during stages N2 and N3 and in association with high-amplitude slow waves, while epileptic activity in turn disturbs sleep^1–6^. During these stages, cortical slow oscillations coordinate thalamocortical spindles and hippocampal ripples, a hierarchy implicated in memory consolidation^7^ and targeted by closed-loop stimulation that enhances slow oscillations and memory^8–11^. Recognizing N2+N3 is therefore central both to studying sleep in epilepsy and to interventions timed to these rhythms.

Sleep is conventionally scored from scalp EEG, electro-oculography (EOG) and chin electromyography (EMG)^12^, although N2 and N3 are primarily defined by EEG features and can be identified from scalp EEG alone^12,13^ In people undergoing stereo-EEG (SEEG), these signals may be incomplete or unreliable: scalp EEG is often absent or non-standard because of SEEG implantation, and the mastoid reference may be contaminated by interictal discharges in temporal lobe epilepsy^14^. Chronically implanted intracranial sensing devices typically record neural activity without concurrent scalp EEG. Scoring from intracranial EEG alone addresses this. The SleepSEEG algorithm, introduced by von Ellenrieder et al., classifies sleep into five stages by first scoring individual intracranial EEG channels and then averaging their predictions across a montage of approximately 95 channels^14^. Developed on recordings from the frontal, parietal, insular, temporal and occipital lobes but not thalamus, it requires a full night of data and does not use anatomical information. Earlier work detected slow-wave sleep^15^ or classified wakefulness, N2 and N3 from intracranial recordings^16^, while single-channel hippocampal recordings enabled classification of wakefulness, REM and NREM in people receiving anterior thalamic stimulation^17^. However, sleep patterns vary across brain structures^18^, so recording site may affect informational content about sleep stage.

The thalamus may be particularly informative for sleep staging, given its role in spindle generation, its involvement in slow oscillation dynamics, and its earlier deactivation relative to the cortex at sleep onset^19–23^. Thalamic leads are increasingly used for epilepsy neuromodulation, particularly in anterior nucleus, with the centromedian and pulvinar nuclei under investigation as alternative targets^24^. Recordings from these nuclei also exhibit day–night differences in spectral activity^25,26^. Whether thalamic channels identify scalp-scored N2+N3 better than other simultaneously recorded intracranial sites, and whether a single electrode suffices for sleep stage classification, remain unclear.

Given this physiology, we hypothesized that thalamic channels would carry more information about N2+N3 than other intracranial sites. We tested this using SEEG recordings with simultaneous scalp EEG from six adults with thalamic contacts, with scalp-scored N2+N3 versus all other stages as the feature of interest. Because a model trained on this cohort could exploit site-specific features, whereas a pretrained model never exposed to thalamic signals could be biased against them, we applied two complementary classifiers to identical bipolar channels: a random forest with hidden Markov decoding (RF + HMM) trained leave-one-participant-out on this cohort, and the pretrained SleepSEEG algorithm. We compared the thalamus with channel-count-matched subsets of other regions and with a white-matter control, and evaluated single bipolar channels and individual thalamic electrodes as representative of what sensing devices record.

## 2 Materials and Methods

### 2.1 Participants and Recordings

Adults with drug-resistant focal epilepsy undergoing SEEG at the University of Chicago were recorded with simultaneous scalp EEG (19–24 scalp electrodes as able based on SEEG electrode location, using the standard 10–20 system). Signals were acquired at 1024–2048 Hz, referenced online to FCz, and downsampled to 256 Hz. Of 31 days of recordings from six participants screened, 25 days met the inclusion criteria of more than 60 minutes of artifact- and seizure-free N2+N3 sleep; days with substantial artifacts or missing scalp EEG were excluded. Each of the 25 days included nocturnal sleep (30–138 h of analyzed data per participant; Supplementary Table S1). The study was approved by the University of Chicago Institutional Review Board (protocol 24-1692), and all participants provided written informed consent.

### 2.2 Sleep Scoring

Sleep staging was performed manually in 30-second epochs by a board-certified clinical sleep physician (K.L.) as wake (W), N1, N2, N3 or REM following AASM criteria^12^, using scalp EEG only because EOG and chin EMG were not recorded. N2 and N3 can be identified relatively reliably from scalp EEG through their characteristic rhythms, whereas REM scoring is less certain without EOG and chin EMG. Together with the thalamocortical involvement in N2 and N3 rhythms, this motivated the selection of N2+N3 as the primary endpoint. Scored, artifact- and seizure-free epochs entered all analyses: 57,199 epochs (477 h), of which 70.4% were W, 1.8% N1, 18.9% N2, 5.3% N3 and 3.5% REM. Every metric was computed in two windows: the whole recording (primary analysis) and the nocturnal sleep period (night).

### 2.3 Anatomical Labeling

Intracranial signals were converted to bipolar derivations between adjacent contacts (bipolar channels), consistent with the montage used by SleepSEEG, after excluding bad contacts and those located in extradural spaces or cerebrospinal fluid. Thalamic SEEG contacts were localized to individual nuclei by co-registering post-implantation CT with pre-implantation MRI, normalizing the images to Montreal Neurological Institute (MNI) space using Lead-DBS, and comparing contact locations with the Morel thalamic atlas^27,28^. Gray-matter contacts were anatomically labeled using the AAL3 atlas^29^ and classified as within the seizure-onset zone (SOZ) or outside (non-SOZ) based on clinical localization, while contacts located in white matter were labeled accordingly. Data from both hemispheres were pooled. The dataset comprised 791 intracranial bipolar channels: 46 thalamic channels across nine nuclei, 106 white-matter channels, 596 channels in 38 AAL3 structures grouped into nine further anatomical systems, and 43 channels spanning two systems or without an AAL3 label (Supplementary Table S2).

### 2.4 Classifiers

#### Random forest with hidden Markov decoding (RF + HMM)

For every channel and epoch, we computed 108 features: 56 broadband, spectral (e.g., aperiodic component), and complexity measures (e.g., Lempel–Ziv complexity^30^), 28 pairwise band-power ratios, and 24 wavelet and multifractal features from SleepSEEG^14^. Features were normalized within each channel and recording. Separate scalp and intracranial random-forest models were trained on class-balanced channel-epochs using leave-one-participant-out cross-validation, with anatomical-system indicators included in the intracranial model. For each test participant, channel outputs were averaged within each evaluation unit and decoded using a hidden Markov model (HMM) with transition probabilities estimated from the training participants (Supplementary Methods S1).

#### SleepSEEG

The pretrained SleepSEEG algorithm^14^, developed without thalamic channels, was applied without retraining. Its features were computed from the same 256-Hz bipolar signals and aligned with the expert-scored epochs. Following SleepSEEG, features were normalized over the night, channel-level probabilities were averaged with equal weight, and sleep stages were assigned using the published decision rule. Alternative normalization windows and the original SleepSEEG processing pipeline gave similar N2+N3 discrimination (Supplementary Methods S2).

#### External scalp scorers

As an external benchmark, pretrained sleep-staging models available through the SLEEPYLAND platform^31^ were applied without retraining to FCz-referenced scalp EEG: U-Sleep^32^, DeepResNet^33^, SleepTransformer^34^, and their ensemble. Sleep-stage labels were determined by majority vote across channels, and the corresponding stage probabilities were used to calculate the area under the receiver operating characteristic curve (AUC; Supplementary Methods S3).

Neither classifier used separate models for individual anatomical regions; evaluation units differed only in the channels averaged at test time, although the random forest included anatomical-system indicators as predictors.

### 2.5 Evaluation Units

Evaluation units included scalp EEG; all intracranial bipolar channels; anatomical systems; AAL3 structures; thalamic nucleus families (pulvinar: medial, lateral, and anterior pulvinar; intralaminar: centromedian and central lateral nuclei; lateral: ventral anterior, ventral lateral and ventral posterolateral nuclei) and individual nuclei; the white-matter control; scalp + thalamus fusion; single bipolar channels; and individual thalamic SEEG electrodes (11 electrodes, 1–2 per participant, each containing 2–6 consecutive thalamic channels). To control for differences in channel count, we sampled up to 30 random subsets of k bipolar channels (k = 1, 2, 3, 4, 6, 8, 12, or each participant’s thalamic channel count) from the all-intracranial, white-matter, mesial temporal, and temporal neocortical pools within each participant, and from thalamic channels for k = 1, 2, 3, 4 and 6 where the participant had at least k thalamic channels, for both classifiers. Single-channel, electrode-level, and subset analyses used the random forest retrained with identical training data, settings and random seeds.

### 2.6 Endpoints and Statistics

The primary endpoint was N2+N3 detection against all other sleep stages, assessed using AUC from stage probabilities, and sensitivity, precision, and F1 from decoded labels. False N2+N3 detections were categorized by expert-scored stage, and precision and coverage of expert-scored N2+N3 were evaluated at probability thresholds of 0.5–0.95. Secondary endpoints included five-stage Cohen’s κ and per-stage F1, excluding stages absent from a recording. The participant was the unit of inference. Recording-level metrics were averaged within participants and summarized as means with bootstrap 95% confidence intervals (CIs; 2,000 resamples). Paired comparisons were reported as participant-level mean differences, their 95% CIs, and the number of participants showing a positive difference. Recording-level linear mixed-effects models with participant-specific random intercepts provided the significance tests, and Benjamini–Hochberg false-discovery-rate correction was applied within each comparison family, analysis window, and metric, with *q* denoting the adjusted mixed-model *p* value. The comparison of the thalamus with mesial temporal channels outside the SOZ used Wilcoxon signed-rank tests on participant means. Units represented in fewer than three participants were excluded from statistical testing. Analyses were performed in MATLAB R2025b.

## 3 Results

### 3.1 Cohort and Study Framework

Figure 1 summarizes the analysis pipeline (Figure 1A), the intracranial coverage (Figure 1B) and an example of expert scoring (Figure 1C). All six participants had thalamic channels (5–11 per participant, 46 in total), as well as mesial temporal, temporal neocortical and white-matter channels (Figure 1B). On the full intracranial montage, the two classifiers reached similar five-stage agreement with the expert over the whole recording (κ 0.57 for RF + HMM and 0.58 for SleepSEEG) and at night (0.40 and 0.44) but made different per-stage errors across anatomical units (Supplementary Tables S3–S5; per-stage F1 and confusion matrices in Supplementary Figures S1–S2).

**Figure 1.**
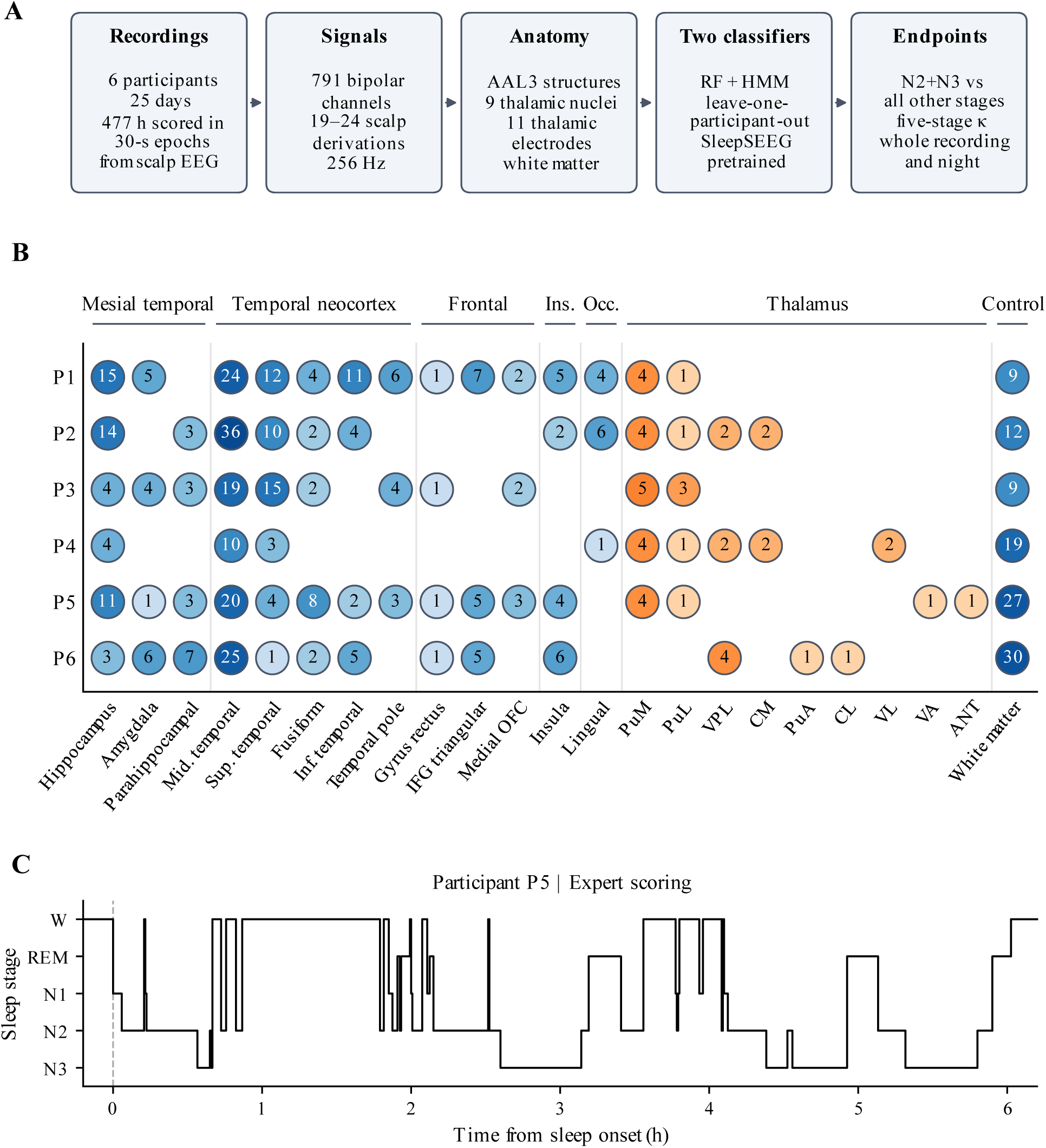
Cohort, intracranial coverage and an example hypnogram. **(A)** Analysis pipeline. Six participants contributed 25 days of recordings (477 h) with simultaneous scalp EEG and stereo-EEG; 791 bipolar channels were labeled anatomically and scored by two classifiers on identical channels. **(B)** Number of bipolar channels per participant (number in each circle; darker shading indicates more channels) for each gray-matter structure present in at least three participants, each thalamic nucleus (orange, any number of participants) and the white-matter control, grouped by anatomical system (Ins., insula; Occ., occipital). PuM, PuL and PuA, medial, lateral and anterior pulvinar; VPL, ventral posterolateral; CM, centromedian; CL, central lateral; VL, ventral lateral; VA, ventral anterior; ANT, anterior nucleus. **(C)** Expert scoring of one night (participant P5), shown from 0.2 h before sleep onset (dashed line). Each 30-s epoch was scored from scalp EEG as W, REM, N1, N2 or N3; the primary endpoint is detection of N2+N3 against all other stages.

### 3.2 Thalamic Channels Detected N2+N3 Better than Temporal and White-Matter Channels

In an example recording, labels derived from the thalamic channels alone followed the expert’s N2+N3 periods with both classifiers (AUC 0.98 and 0.97) despite moderate five-stage agreement (Figure 2A). Across participants, detection of N2+N3 was highest on scalp (AUC 0.96) and, among the systems sampled in all six participants, on the thalamus (0.94 with RF + HMM, 0.93 with SleepSEEG; Figure 2B, Table 1). With RF + HMM, the thalamus was close to scalp (difference −0.015, 95% CI −0.037 to 0.004, q = 0.16) and exceeded the mesial temporal lobe (+0.068, 0.027 to 0.111, six of six participants, q = 0.024), the hippocampus (+0.070, six of six, q = 0.024) and the temporal neocortex (+0.064, six of six, q = 0.024); it was higher than the full intracranial montage in five of six participants (+0.032, 0.007 to 0.067, q = 0.064; Supplementary Table S6). With SleepSEEG, thalamic AUC was higher than that of the full montage, mesial temporal lobe, hippocampus, temporal neocortex and white matter (+0.024 to +0.073) but not significantly after correction (q ≥ 0.08). Each thalamic nucleus and nucleus family tested reached 0.88–0.95 with RF + HMM and 0.86–0.92 with SleepSEEG (Table 1). The same ranking held for F1 of the decoded labels: thalamus 0.74 vs mesial temporal 0.62 (+0.11, six of six, q = 0.008) with RF + HMM, and with SleepSEEG the thalamus had the highest F1 of the regions compared (0.78 vs 0.69–0.73; vs temporal neocortex +0.087, six of six, q = 0.004; Supplementary Figure S3). At night, thalamic AUC matched that of RF + HMM on scalp (0.88 vs 0.88) and was above the mesial temporal lobe (+0.061, six of six, q = 0.018). Adding the thalamus to scalp increased AUC slightly but consistently (0.966 vs 0.958, +0.008, five of six, q = 0.011). For five-stage scoring the thalamus did not stand out: its κ with RF + HMM (0.52) was lower than scalp in all six participants (q = 0.013) and similar to other intracranial units, because thalamic signals separated N2 from N3 poorly (N3 F1 0.27 vs 0.42–0.48 for scalp, the full montage, the mesial temporal lobe and white matter; q ≤ 0.004) and REM poorly (F1 0.28).

**Figure 2.**
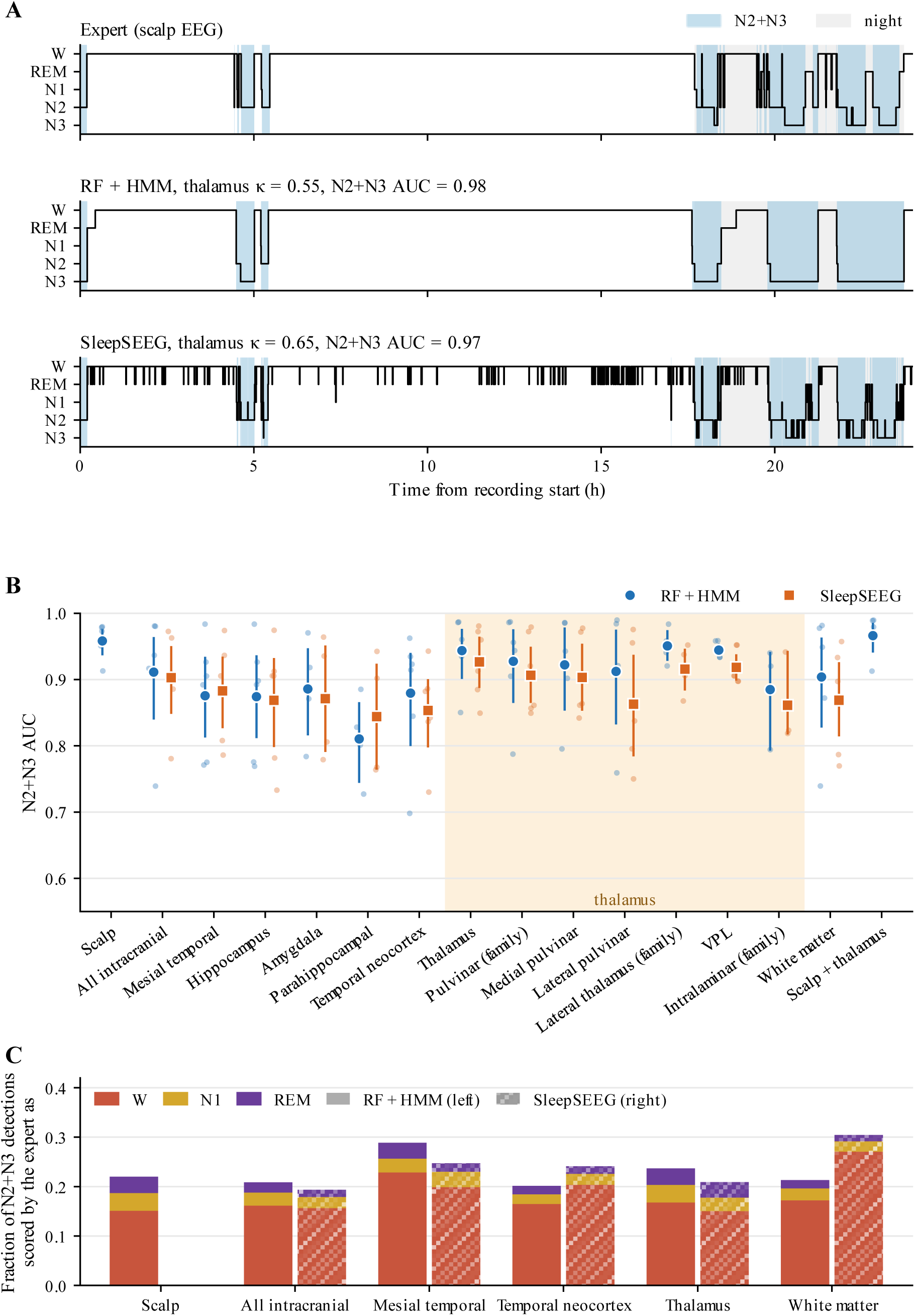
Detection of N2+N3 sleep across regions, whole recording. **(A)** The recording in Figure 1C over the whole 24 h: expert scoring and the labels produced from the thalamic channels alone by RF + HMM and by SleepSEEG. Blue, epochs labeled N2+N3; gray, night. κ, five-stage Cohen’s kappa; AUC for N2+N3 versus all other stages, for this recording. Both classifiers separated N2+N3 from the other stages; five-stage agreement was lower mainly because RF + HMM labeled most of the expert’s N2 and REM epochs as N3, and SleepSEEG labeled daytime wake as REM. **(B)** N2+N3 AUC by unit for RF + HMM (circles) and SleepSEEG (squares): scalp, the full intracranial montage, the systems sampled in all six participants with their structures, nuclei or nucleus families sampled in at least three participants, and scalp + thalamus fusion. Small dots are participants (recordings averaged within participant); markers and bars are the mean and bootstrap 95% confidence interval across participants. Shading marks thalamic units; SleepSEEG is not defined for scalp or scalp + thalamus fusion. **(C)** Composition of false N2+N3 detections: fraction of the epochs each classifier labeled N2+N3 that the expert scored as W, N1 or REM (left bars, RF + HMM; right, hatched bars, SleepSEEG), mean across participants. Bar height equals one minus precision.

**Table 1.** Detection of N2+N3 sleep and five-stage agreement by unit. Values are means across participants for RF + HMM / SleepSEEG over the whole recording (Whole) and at night (Night); n, number of participants in whom the unit was sampled. Rows are grouped by system: the scalp reference, the full intracranial montage, the anatomical systems sampled in all six participants (mesial temporal lobe, temporal neocortex, thalamus and the white-matter control) and scalp + thalamus fusion. Within the mesial temporal lobe and the thalamus, the first row pools all channels of the system and the rows below it are the structures or nuclei sampled in at least three participants. AUC for N2+N3 versus all other stages; F1 of the decoded N2+N3 labels; κ, five-stage Cohen’s kappa; “–”, not defined (SleepSEEG does not score scalp).

| **System** | **Unit** | **n** | **N2+N3 AUC** |  | **N2+N3 F1** |  | **Five-stage κ** |  |
|---|---|---|---|---|---|---|---|---|
|  |  |  | **Whole** | **Night** | **Whole** | **Night** | **Whole** | **Night** |
| Scalp | Scalp | 6 | 0.96 / – | 0.88 / – | 0.72 / – | 0.74 / – | 0.59 / – | 0.43 / – |
| All intracranial channels | All intracranial channels | 6 | 0.91 / 0.90 | 0.85 / 0.82 | 0.72 / 0.78 | 0.73 / 0.80 | 0.57 / 0.58 | 0.40 / 0.44 |
| Mesial temporal lobe | All mesial temporal channels | 6 | 0.88 / 0.88 | 0.82 / 0.81 | 0.62 / 0.73 | 0.64 / 0.76 | 0.47 / 0.51 | 0.32 / 0.41 |
|  | Hippocampus | 6 | 0.87 / 0.87 | 0.81 / 0.80 | 0.61 / 0.69 | 0.64 / 0.73 | 0.46 / 0.45 | 0.30 / 0.36 |
|  | Amygdala | 4 | 0.89 / 0.87 | 0.85 / 0.85 | 0.66 / 0.75 | 0.68 / 0.80 | 0.48 / 0.45 | 0.35 / 0.43 |
|  | Parahippocampal gyrus | 4 | 0.81 / 0.84 | 0.76 / 0.78 | 0.52 / 0.63 | 0.57 / 0.70 | 0.36 / 0.34 | 0.28 / 0.31 |
| Temporal neocortex | All temporal neocortical channels | 6 | 0.88 / 0.85 | 0.81 / 0.78 | 0.65 / 0.70 | 0.65 / 0.73 | 0.50 / 0.48 | 0.31 / 0.36 |
| Thalamus | All thalamic channels | 6 | 0.94 / 0.93 | 0.88 / 0.86 | 0.74 / 0.78 | 0.76 / 0.80 | 0.52 / 0.42 | 0.31 / 0.38 |
|  | Pulvinar family (PuM, PuL, PuA) | 6 | 0.93 / 0.91 | 0.87 / 0.83 | 0.73 / 0.75 | 0.75 / 0.78 | 0.49 / 0.38 | 0.28 / 0.33 |
|  | Medial pulvinar | 5 | 0.92 / 0.90 | 0.86 / 0.85 | 0.71 / 0.76 | 0.73 / 0.79 | 0.47 / 0.39 | 0.29 / 0.33 |
|  | Lateral pulvinar | 5 | 0.91 / 0.86 | 0.86 / 0.81 | 0.73 / 0.68 | 0.76 / 0.75 | 0.51 / 0.25 | 0.33 / 0.27 |
|  | Ventral posterolateral | 3 | 0.94 / 0.92 | 0.84 / 0.82 | 0.71 / 0.68 | 0.74 / 0.72 | 0.50 / 0.28 | 0.25 / 0.22 |
|  | Intralaminar (CM, CL) | 3 | 0.88 / 0.86 | 0.81 / 0.79 | 0.68 / 0.62 | 0.71 / 0.69 | 0.41 / 0.20 | 0.24 / 0.21 |
|  | Lateral thalamus family (VA, VL, VPL) | 4 | 0.95 / 0.92 | 0.87 / 0.85 | 0.75 / 0.70 | 0.77 / 0.75 | 0.54 / 0.28 | 0.35 / 0.28 |
| White matter (control) | White-matter control | 6 | 0.90 / 0.87 | 0.84 / 0.80 | 0.72 / 0.69 | 0.75 / 0.76 | 0.56 / 0.40 | 0.41 / 0.38 |
| Fusion | Scalp + thalamus | 6 | 0.97 / – | 0.89 / – | 0.73 / – | 0.73 / – | 0.62 / – | 0.46 / – |

Because the thalamus was compared with RF + HMM on scalp, we benchmarked that model against pretrained scalp scorers applied to the same recordings (Supplementary Table S7, Supplementary Figure S4). The three deep-learning networks and their ensemble reached an N2+N3 AUC of 0.96–0.97 over the whole recording, equal to RF + HMM on scalp (differences ≤ 0.01, q ≥ 0.39), with higher five-stage agreement (κ 0.73–0.76 vs 0.59) and N2+N3 F1 (0.86–0.88 vs 0.72). At night their N2+N3 AUC (0.91–0.92) was higher than that of RF + HMM on scalp in all six participants (+0.031 to +0.037, q = 0.087) and above that of the thalamus (0.88).

### 3.3 The Thalamic Advantage Withstands Control Analyses

Detection improved with the number of channels averaged, and participants had more mesial temporal or white-matter channels than thalamic channels, so we compared the thalamus with random subsets of other channels (Figure 3A). When subsampled in the same way, the thalamus had the highest mean AUC at every k tested (1, 2, 3, 4 and 6) with both classifiers, and with RF + HMM a single thalamic channel (0.92) matched or exceeded twelve channels from any of the four comparison pools (≤ 0.91). Matched to each participant’s own number of thalamic channels (5–11), the thalamus exceeded subsets of all intracranial channels (+0.036, 0.013 to 0.063, six of six participants), white matter (+0.043, 0.015 to 0.075, five of six), mesial temporal channels (+0.082, 0.035 to 0.131, five of five) and temporal neocortical channels (+0.069, 0.033 to 0.106, six of six) with RF + HMM (q = 0.011 for each), and the same held at night (+0.036 to +0.074, q ≤ 0.030; Supplementary Table S8). With SleepSEEG, the thalamus exceeded matched white-matter (AUC +0.068, five of six, q = 0.029) and temporal neocortical subsets (+0.092, six of six, q = 0.018), and its N2+N3 F1 exceeded matched white-matter (+0.111, six of six, q < 0.001), temporal neocortical (+0.123, six of six, q < 0.001) and mesial temporal subsets (+0.090, four of five, q = 0.020).

**Figure 3.**
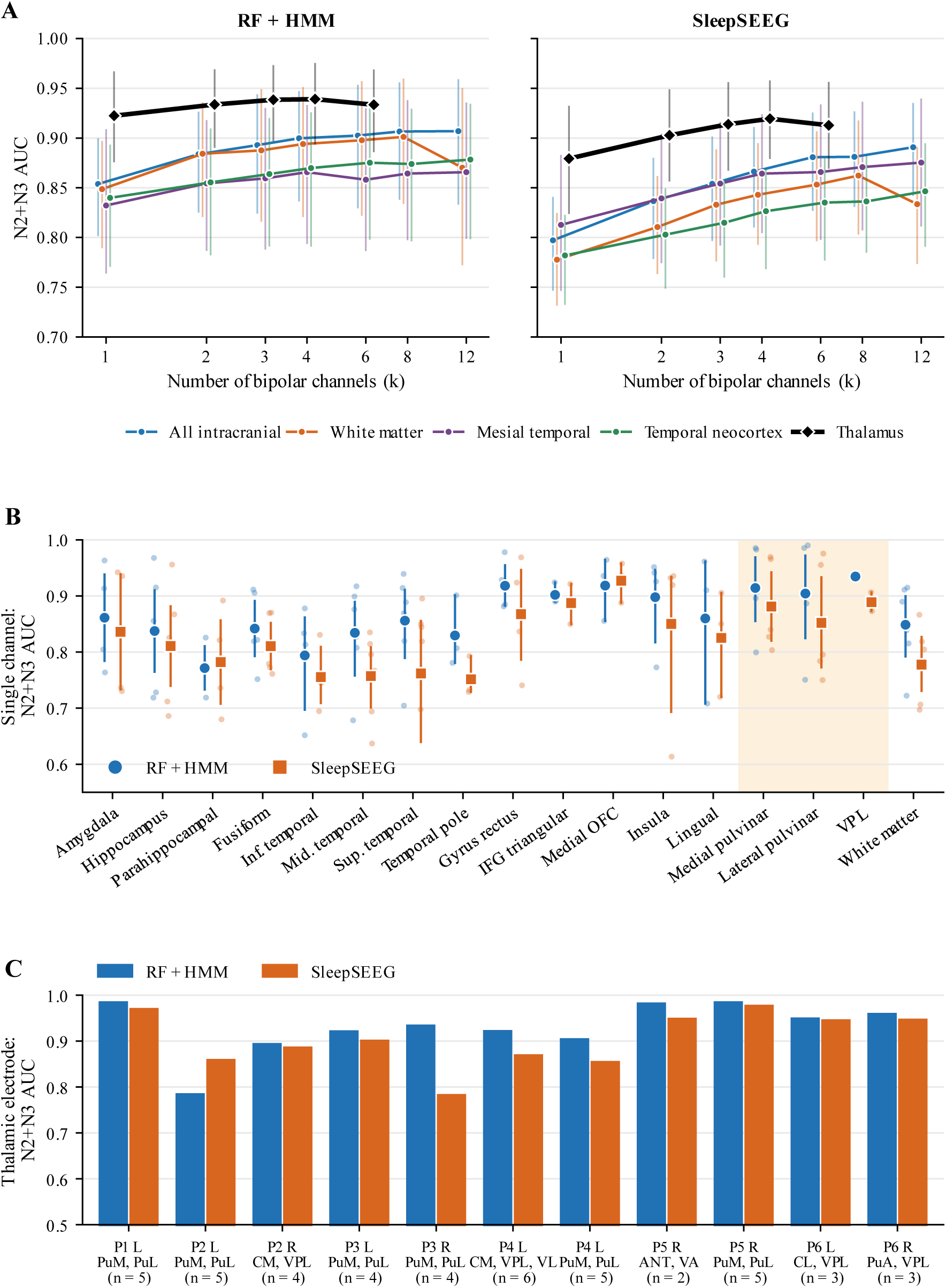
Channel-count control and detection from single channels and single electrodes, whole recording. **(A)** N2+N3 AUC when the outputs of k randomly chosen bipolar channels are averaged (up to 30 subsets per k and recording), drawn from all intracranial, white-matter, mesial temporal and temporal neocortical channels and from the thalamus (black), with RF + HMM (left) and SleepSEEG (right). Thalamic subsets were drawn for k = 1, 2, 3, 4 and 6 in participants with at least k thalamic channels (all six participants for k ≤ 4, five for k = 6). Points are means across participants with bootstrap 95% CI, shown where at least three participants contributed; logarithmic x-axis. **(B)** N2+N3 AUC of single bipolar channels, each scored on its own, by structure (structures with channels in at least three participants; thalamic nuclei shaded). Small dots are participants (mean over that participant’s channels in the structure); markers and bars are the mean and bootstrap 95% CI across participants. **(C)** N2+N3 AUC of each thalamic electrode, scored with all thalamic channels on that electrode; labels give the participant, hemisphere, the nuclei traversed from the deepest contact outward and the number of thalamic channels (n). The night version is Supplementary Figure S6.

White-matter channels mainly pick up activity volume-conducted from surrounding gray matter and therefore test whether a region carries more stage information than spatially diffuse signals. The thalamus exceeded the white-matter control for N2+N3 with both classifiers (RF + HMM AUC +0.040, five of six, q = 0.036; SleepSEEG F1 +0.092, six of six, q = 0.002; Supplementary Table S6). How other systems ranked against white matter for five-stage scoring, by contrast, depended on the classifier, and no other system differed from white matter in N2+N3 AUC or F1 with either classifier (Supplementary Table S9), so the thalamic N2+N3 advantage was the only system-level advantage over white matter consistent across classifiers.

Three further analyses addressed explanations unrelated to thalamic signals. SleepSEEG, developed without thalamic channels, also ranked the thalamus highest of the systems sampled in all six participants (Table 1), so the ranking did not depend on a model trained on this cohort’s thalamic data. In the random forest, anatomical-system indicators accounted for 0.6% of intracranial predictor importance, whereas Lempel–Ziv complexity and zero-crossing rate ranked highest (Supplementary Figure S5). Removing SOZ channels did not improve N2+N3 detection on the mesial temporal lobe (AUC −0.006 with RF + HMM, −0.030 with SleepSEEG), and the thalamus still exceeded mesial temporal channels outside the SOZ (RF + HMM +0.074, six of six; SleepSEEG +0.073, five of six; Supplementary Table S10).

### 3.4 Single Channels, Single Electrodes and Detection Errors

Over the whole recording, a single thalamic bipolar channel detected N2+N3 with AUC 0.90–0.93 with RF + HMM (ventral posterolateral 0.93, medial pulvinar 0.91, lateral pulvinar 0.90), compared with 0.77–0.86 for single mesial temporal and temporal neocortical channels and 0.85 for single white-matter channels (Figure 3B). Single channels in the medial orbitofrontal cortex, gyrus rectus and the triangular part of the inferior frontal gyrus, sampled in three or four participants, performed comparably (0.90–0.92). Every thalamic electrode reached AUC 0.79–0.99 with RF + HMM (10 of 11 ≥ 0.89) and 0.79–0.98 with SleepSEEG. This included the two electrodes entering the centromedian nucleus (0.90 and 0.92) and the single anterior-nucleus electrode (0.98; Figure 3C), although each of these clinical targets was sampled in only one or two participants.

Most false N2+N3 detections were epochs scored as wake. On the thalamus, 17% of the N2+N3 detections of RF + HMM were expert wake, 4% N1 and 3% REM over the whole recording, and 15%, 3% and 4% at night; other units showed the same pattern (Figure 2C, Supplementary Figure S3). Accepting only high-confidence detections (N2+N3 probability ≥ 0.95 instead of ≥ 0.5) reduced false detections. Over the whole recording, RF + HMM still identified 72% of the expert’s N2+N3 on the thalamus, 63% on scalp and 51% on the mesial temporal lobe, and 82%, 86% and 79% of its detections, respectively, were correct.

## 4 Discussion

In this study, thalamic channels recognized scalp-scored N2+N3 sleep better than the other intracranial regions sampled in every participant, as hypothesized from thalamic physiology. Their N2+N3 discrimination approached that of scalp EEG, exceeded the mesial temporal lobe and temporal neocortex in every participant with RF + HMM over the whole recording, and held against channel-count-matched white-matter and temporal neocortical subsets with both classifiers in both analysis windows. A single thalamic electrode retained most of this discrimination. The thalamus separated N2 from N3 poorly, which an N2+N3 detector does not require.

The pattern fits the role of the thalamus in NREM sleep. The thalamus contributes to spindle generation and slow oscillation dynamics, with anterior thalamic events preceding their neocortical counterparts^19^. Thalamic signals therefore change markedly at the onset of NREM sleep, separating N2+N3 from wake and REM. Because spindles, the defining scalp marker of N2, are thalamocortical events, close agreement with scalp-scored N2+N3 is expected. The N2/N3 boundary, by contrast, is defined by the amount of high-amplitude slow waves on scalp EEG, a cortical property that thalamic recordings track less directly. Slow oscillations and spindles occur during both N2 and N3, when epileptiform activity is also commonly enhanced, making an N2+N3 detector relevant for monitoring and stimulation timed to these rhythms.

The two classifiers reached similar overall agreement but made different errors, partly reflecting training data. The random forest was trained on this study’s cohort, thalamus included, whereas SleepSEEG classified wake and REM less well, often labeling wake as REM, on thalamic and white-matter channels, which lie outside its training distribution (Supplementary Table S3, Supplementary Figure S2). Because the classifier also changed how regions ranked against white matter for five-stage scoring, a regional claim is credible only when it holds under both models, as the thalamic N2+N3 advantage did.

As a check on the scalp reference, pretrained scalp scorers labeled the five stages better than RF + HMM on scalp, with agreement close to that between human expert scorers (κ 0.76)^35^, but did not discriminate N2+N3 better over the whole recording, so the scalp reference was not weak for the primary endpoint. Their decoded N2+N3 labels were, however, more accurate, and at night their N2+N3 AUC was higher in all six participants, although not significantly after correction, so the thalamus approaches, rather than matches, the best scalp scoring.

For sleep monitoring in people with implanted devices and for closed-loop neuromodulation, these results identify which channels carry the information a stage detector needs; they do not show that a device can use it in real time. Both classifiers normalized features over the recording or the night, and the hidden Markov model decoded the whole sequence, using information a stimulator would not have when deciding to deliver a stimulus, so the reported performance is an upper bound. Within that bound, three findings bear on device design. Thalamic channels alone approach scalp-level N2+N3 discrimination. The residual errors are mainly wake epochs, where false N2+N3 detections could lead to unintended stimulation, and they decrease when only high-confidence detections are accepted. Adding scalp to thalamic signals improves detection slightly but consistently. Causal, epoch-by-epoch evaluation using thalamic channels is the necessary next step.

This study has limitations. The cohort was small and derived from a single center; consistency across participants and classifiers supports the main result. Sleep stages were scored from scalp EEG without EOG and EMG. Results therefore describe agreement with scalp-scored stages rather than full polysomnography, and REM labels are the least reliable. Because the expert scored the same scalp signals that the scalp classifiers used, the comparison favors scalp over intracranial channels. Only the temporal lobe, thalamus and white matter were sampled in every participant; frontal-lobe channels, sampled in five participants, performed comparably to the thalamus with RF + HMM (AUC 0.93 vs 0.94), and cingulate channels, sampled in three, reached 0.96. Thalamic placement was clinically determined and heterogeneous: nuclei within a participant lay along one or two electrodes and were not independent samples, and centromedian and anterior-nucleus results rest on one or two participants. The random forest received a coarse anatomical indicator, but SleepSEEG, which uses no anatomical information, gave the same ranking. SEEG recordings are dominated by wake, which inflates the weight of wake in five-stage κ; we therefore also report results at night. Finally, all participants had epilepsy and were monitored in the hospital, where sleep is disrupted.

In conclusion, thalamic SEEG identifies scalp-scored N2+N3 sleep with discrimination approaching that of scalp EEG and better than the temporal-lobe and white-matter channels sampled in every participant, down to a single electrode and with two complementary classifiers. Because the thalamus is already a target for stimulation in epilepsy, these offline results suggest that the same leads could also sense sleep state, supporting thalamic sleep monitoring and the development of causal thalamic N2+N3 detection for sleep-targeted neuromodulation.

## Data availability statement

The data supporting the findings of this study may be made available upon reasonable request, in accordance with institutional policies and ethical requirements for sharing human participant data.

## Ethics statement

The study involving humans was approved by the University of Chicago Institutional Review Board (protocol 24-1692). Participants provided written informed consent.

## References

1. Frauscher, B. et al. Facilitation of epileptic activity during sleep is mediated by high amplitude slow waves. Brain 138, 1629–1641 (2015).

2. Frauscher, B. & Gotman, J. Sleep, oscillations, interictal discharges, and seizures in human focal epilepsy. Neurobiology of disease 127, 545–553 (2019).

3. Ng, M. & Pavlova, M. Why are seizures rare in rapid eye movement sleep? Review of the frequency of seizures in different sleep stages. Epilepsy research and treatment 2013, 932790 (2013).

4. Peter-Derex, L., et al. Sleep disruption in epilepsy: ictal and interictal epileptic activity matter. Annals of Neurology 88, 907–920 (2020).

5. Fouad, A. et al. Interictal epileptiform discharges show distinct spatiotemporal and morphological patterns across wake and sleep. Brain Communications 4, fcac183 (2022).

6. Blasco, X. et al. Sleep stage-dependent distribution of interictal epileptiform discharges in epilepsy: A systematic review. Epilepsy & Behavior 184, 111233 (2026).

7. Staresina, B. P., Niediek, J., Borger, V., Surges, R. & Mormann, F. How coupled slow oscillations, spindles and ripples coordinate neuronal processing and communication during human sleep. Nat Neurosci 26, 1429–1437 (2023).

8. Ngo, H.-V. V., Martinetz, T., Born, J. & Mölle, M. Auditory closed-loop stimulation of the sleep slow oscillation enhances memory. Neuron 78, 545–553 (2013).

9. Geva-Sagiv, M. et al. Augmenting hippocampal–prefrontal neuronal synchrony during sleep enhances memory consolidation in humans. Nature neuroscience 26, 1100–1110 (2023).

10. Alipour, M., van Drongelen, W., Malerba, P., Voss, J. & Satzer, D. Predictability of sleep slow oscillation emergence and spatial extent from pre-onset neural dynamics. Scientific Reports (2026).

11. Alipour, M., Seok, S., Mednick, S. C. & Malerba, P. A classification-based generative approach to selective targeting of global slow oscillations during sleep. Frontiers in Human Neuroscience 18, 1342975 (2024).

12. Berry, R. B. et al. AASM scoring manual updates for 2017 (version 2.4). Journal of clinical sleep medicine 13, 665–666 (2017).

13. Lucey, B. P. et al. Comparison of a single-channel EEG sleep study to polysomnography. Journal of sleep research 25, 625–635 (2016).

14. von Ellenrieder, N., Peter-Derex, L., Gotman, J. & Frauscher, B. SleepSEEG: automatic sleep scoring using intracranial EEG recordings only. Journal of Neural Engineering 19, 026057 (2022).

15. Reed, C. M. et al. Automatic detection of periods of slow wave sleep based on intracranial depth electrode recordings. Journal of Neuroscience Methods 282, 1–8 (2017).

16. Kremen, V. et al. Automated unsupervised behavioral state classification using intracranial electrophysiology. J. Neural Eng. 16, 026004 (2019).

17. Mivalt, F. et al. Electrical brain stimulation and continuous behavioral state tracking in ambulatory humans. J. Neural Eng. 19, 016019 (2022).

18. Von Ellenrieder, N. et al. How the Human Brain Sleeps: Direct Cortical Recordings of Normal Brain Activity. Annals of Neurology 87, 289–301 (2020).

19. Schreiner, T., Kaufmann, E., Noachtar, S., Mehrkens, J.-H. & Staudigl, T. The human thalamus orchestrates neocortical oscillations during NREM sleep. Nature communications 13, 5231 (2022).

20. Gent, T. C., Bandarabadi, M., Herrera, C. G. & Adamantidis, A. R. Thalamic dual control of sleep and wakefulness. Nature neuroscience 21, 974–984 (2018).

21. Magnin, M., et al. Thalamic deactivation at sleep onset precedes that of the cerebral cortex in humans. Proc. Natl. Acad. Sci. U.S.A. 107, 3829–3833 (2010).

22. Alipour, M., Mednick, S. C. & Malerba, P. Sleep slow oscillation emergence on the scalp as a renewal point process. PLOS Computational Biology 22, e1014572 (2026).

23. Alipour, M., Drongelen, W. van, Jacobs, J., Voss, J. & Satzer, D. Thalamic Nuclei Differentially Coordinate Propagation of Cortical Slow Oscillations. bioRxiv 2026–06 (2026).

24. Hsu, J. et al. Thalamic Neuromodulation: New Frontiers. Epilepsy Curr 15357597261416388 (2026) doi:10.1177/15357597261416388.

25. Satzer, D. et al. Ambulatory Local Field Potential Recordings from the Thalamus in Epilepsy: A Feasibility Study. Stereotact Funct Neurosurg 101, 195–206 (2023).

26. Burdette, D., Patra, S., Morrell, M. & Johnson, L. Clusters in thalamic iEEG spectral features reveal sleep/wake differences over time. Journal of Clinical Neurophysiology 43, 455 (2026).

27. Niemann, K., Mennicken, V., Jeanmonod, D. & Morel, A. The Morel stereotactic atlas of the human thalamus: atlas-to-MR registration of internally consistent canonical model. Neuroimage 12, 601–616 (2000).

28. Horn, A. et al. Lead-DBS v2: Towards a comprehensive pipeline for deep brain stimulation imaging. Neuroimage 184, 293–316 (2019).

29. Rolls, E. T., Huang, C.-C., Lin, C.-P., Feng, J. & Joliot, M. Automated anatomical labelling atlas 3. Neuroimage 206, 116189 (2020).

30. Alipour, M. & Hashemi Gholpayeghani, S. M. R. Real-time non-uniform EEG sampling. Biomedical Signal Processing and Control 70, 102961 (2021).

31. Rossi, A. D. et al. SLEEPYLAND: trust begins with fair evaluation of automatic sleep staging models. npj Digital Medicine 9, 55 (2025).

32. Perslev, M. et al. U-Sleep: resilient high-frequency sleep staging. NPJ digital medicine 4, 72 (2021).

33. Olesen, A. N., Jørgen Jennum, P., Mignot, E. & Sorensen, H. B. D. Automatic sleep stage classification with deep residual networks in a mixed-cohort setting. Sleep 44, zsaa161 (2021).

34. Phan, H. et al. Sleeptransformer: Automatic sleep staging with interpretability and uncertainty quantification. IEEE Transactions on Biomedical Engineering 69, 2456–2467 (2022).

35. Lee, Y. J., Lee, J. Y., Cho, J. H. & Choi, J. H. Interrater reliability of sleep stage scoring: a meta-analysis. Journal of clinical sleep medicine 18, 193–202 (2022).

