## Supplementary Materials for "Thalamic stereo-EEG detects sleep stages N2 and N3 with discrimination approaching scalp EEG"

### Supplementary Material

### Supplementary Methods

#### Supplementary Methods S1. Random-forest preprocessing, features, training, and decoding

For random-forest feature extraction, signals were filtered using zero-phase second-order filters (0.3-Hz high-pass and 58–62 Hz notch) and band-passed into eight frequency bands: slow oscillation (SO, 0.5–2 Hz), delta (2–4 Hz), theta (4–8 Hz), alpha (8–12 Hz), sigma (12–16 Hz), beta (16–30 Hz), gamma (30–58 Hz), and high gamma (62–118 Hz). For each channel and 30-s epoch, 108 features were computed in five families:

- **Broadband (10):** Log variance, Hjorth mobility and complexity, skewness, kurtosis, log line length, zero-crossing rate, permutation entropy (order 3), Lempel–Ziv complexity, and Higuchi fractal dimension. The last two were computed in 2-s blocks and averaged.
- **Spectral (22):** Log and relative power in eight bands, spectral entropy, 95% spectral edge frequency, spectral centroid, aperiodic exponent and offset (power-law fit over 2–40 Hz), and the 30–45 Hz spectral slope.
- **Band-specific (24):** Hjorth mobility, Hjorth complexity, and permutation entropy for each band-passed signal.
- **Band ratios (28):** All pairwise log-power ratios across the eight frequency bands.
- **SleepSEEG (24):** Wavelet and multifractal features from SleepSEEG, computed on the unfiltered bipolar signal using a 32-s window centered on each epoch.

Features were normalized offline within each channel and recording using robust z-scores (subtracting the median and dividing by the interquartile range/1.349), smoothed with a three-epoch moving mean, and clipped at ±8. Separate scalp and intracranial random-forest models were trained using leave-one-participant-out cross-validation. In each fold, training data comprised class-balanced channel-epochs from five participants, with a maximum of 120,000 training samples per model. Each model contained 100 bagged classification trees (minimum leaf size 5; $\sqrt{p}$ randomly selected predictors per split; MATLAB fitcensemble). The intracranial model also received anatomical-system indicators as predictors.

For each evaluation unit, channel-level outputs were averaged with equal weight, following SleepSEEG. Hidden Markov model decoding used stage-transition probabilities and stage priors estimated from the training participants' expert hypnograms. Final sleep-stage labels were obtained using the Viterbi algorithm, while forward–backward posterior probabilities were used to calculate AUC and apply classification thresholds. For scalp + thalamus fusion, scalp and thalamic predictions were combined using the normalized geometric mean of their stage probabilities.

**Supplementary Methods S2. SleepSEEG implementation and robustness checks**

SleepSEEG (the version blind to epileptic activity) was applied without retraining. Each channel was assigned to one of 20 pretrained channel clusters and classified independently, after which channel probabilities were averaged with equal weight and sleep stages assigned using the published decision rule. This allowed SleepSEEG to be evaluated on individual anatomical units by averaging only their corresponding channels. SleepSEEG features were computed from the same 256-Hz bipolar signals used by the random forest, with 30-s epochs aligned to the expert annotations. As a robustness check, we also ran the original SleepSEEG pipeline, which reads the EDF file and defines its own 30-s epochs from the start of the file, to check that aligning the epochs to the expert scoring did not change its performance: on the 18 recordings for which the original EDF files were available, N2+N3 AUC differed by at most 0.01 between the two pipelines over the whole recording (all intracranial channels 0.885 aligned vs 0.882 original; thalamus 0.895 vs 0.885). Features were normalized over the nocturnal sleep period marked on the hypnogram. To exclude any dependence on the expert scoring, SleepSEEG was re-run with two normalization windows independent of it: a fixed clock window (22:00–07:00, from the recording start time) and the whole recording. Thalamic N2+N3 AUC changed by at most 0.008, and the thalamus remained the unit with the highest N2+N3 AUC under every window.

**Supplementary Methods S3. External scalp scorers**

Pretrained scalp-EEG sleep-staging models distributed through SLEEPYLAND were applied without retraining or fine-tuning: U-Sleep, DeepResNet, SleepTransformer and their ensemble. All available FCz-referenced scalp EEG derivations were used, excluding the mastoid channel; EOG and EMG were unavailable. Each model scored individual channels in 30-s epochs, with sleep-stage labels determined by SLEEPYLAND's majority vote across channels. AUC was calculated from the corresponding five-stage probabilities, with N2+N3 probability defined as the sum of N2 and N3 probabilities. Evaluation used the same artifact- and seizure-free epochs, analysis windows, and statistical procedures as the other classifiers, with participants as the unit of inference. Each model was compared with the scalp RF + HMM reference, with false-discovery-rate correction across scorers within each window and metric.

### Supplementary Tables

**Supplementary Table S1. Recording duration, expert-scored sleep-stage epochs, and channel coverage by participant.** Across 25 days, 57,199 artifact- and seizure-free 30-s epochs were scored: 70.4% wake (W), 1.8% N1, 18.9% N2, 5.3% N3, and 3.5% REM. Four days contained no REM sleep, and one contained no N3 sleep. Scalp derivations, number of scalp EEG derivations; SOZ, seizure-onset zone.

| Participant | Recorded data (h) | Artifact- and seizure-free scored data (h) | W epochs | N1 epochs | N2 epochs | N3 epochs | REM epochs | Scalp derivations | Bipolar channels | Thalamic channels | White-matter channels | SOZ channels |
|---|---|---|---|---|---|---|---|---|---|---|---|---|
| P1 | 143 | 137.9 | 11317 | 317 | 3234 | 791 | 891 | 19 | 150 | 5 | 9 | 33 |
| P2 | 95.8 | 93.4 | 9402 | 208 | 1412 | 121 | 60 | 24 | 149 | 9 | 12 | 10 |
| P3 | 60.1 | 59.3 | 4480 | 99 | 1538 | 851 | 153 | 24 | 97 | 8 | 9 | 13 |
| P4 | 92.7 | 65.4 | 5245 | 141 | 1511 | 571 | 378 | 24 | 124 | 11 | 19 | 8 |
| P5 | 30.7 | 30.1 | 2776 | 40 | 476 | 236 | 83 | 24 | 143 | 7 | 27 | 6 |
| P6 | 93.9 | 90.6 | 7066 | 249 | 2653 | 489 | 411 | 24 | 128 | 6 | 30 | 8 |
| Total | 516.2 | 476.7 | 40286 | 1054 | 10824 | 3059 | 1976 | 19–24 | 791 | 46 | 106 | 78 |

**Supplementary Table S2. Anatomical coverage: bipolar channels per participant in each anatomical system, with the AAL3 structures or thalamic nuclei of the system.** System counts include channels spanning two structures of the same system; structures and nuclei are listed with their number of channels. *, tested (sampled in at least three participants). The last two rows give the channels spanning two systems or without an AAL3 label, which entered only the all-intracranial unit, and all intracranial channels. No thalamic channel was in the seizure-onset zone (SOZ); ·, none.

| System | P1 | P2 | P3 | P4 | P5 | P6 | Participants | Channels | SOZ channels | AAL3 structures or thalamic nuclei (channels) |
|---|---|---|---|---|---|---|---|---|---|---|
| Mesial temporal lobe* | 21 | 18 | 14 | 4 | 18 | 22 | 6 | 97 | 38 | Hippocampus* (51); Amygdala* (16); Parahippocampal gyrus* (16) |
| Temporal neocortex* | 72 | 58 | 44 | 14 | 46 | 36 | 6 | 270 | 26 | Middle temporal gyrus* (134); Superior temporal gyrus* (45); Inferior temporal gyrus* (22); Fusiform gyrus* (18); Temporal pole, middle part* (13); Temporal pole, superior part (12); Heschl's gyrus (2) |
| Frontal* | 22 | · | 19 | 4 | 18 | 10 | 5 | 73 | 0 | Inferior frontal gyrus, triangular part* (17); Medial orbitofrontal cortex* (7); Gyrus rectus* (4); Inferior frontal gyrus, orbital part (10); Anterior orbitofrontal cortex (6); Middle frontal gyrus (5); Superior frontal gyrus, dorsolateral (4); Lateral orbitofrontal cortex (3); Medial frontal gyrus, orbital part (3); Posterior orbitofrontal cortex (2) |
| Insula / operculum* | 5 | 2 | · | · | 9 | 6 | 4 | 22 | 0 | Insula* (17); Rolandic operculum (4) |
| Cingulate* | 2 | · | · | 6 | 5 | · | 3 | 13 | 0 | Anterior cingulate cortex, supracallosal (7); Middle cingulate cortex (6) |
| Sensorimotor | · | · | · | 31 | · | 8 | 2 | 39 | 0 | Postcentral gyrus (17); Precentral gyrus (10); Paracentral lobule (8); Supplementary motor area (2) |
| Parietal | · | 33 | · | 16 | · | · | 2 | 49 | 4 | Precuneus (19); Angular gyrus (16); Superior parietal lobule (10); Inferior parietal lobule (1) |
| Occipital* | 4 | 9 | · | 13 | · | · | 3 | 26 | 4 | Lingual gyrus* (11); Cuneus (6); Middle occipital gyrus (3); Calcarine cortex (1); Superior occipital gyrus (1) |
| Basal ganglia | · | · | · | · | 1 | 6 | 2 | 7 | 0 | Putamen (7) |

| System | P1 | P2 | P3 | P4 | P5 | P6 | Participants | Channels | SOZ channels | AAL3 structures or thalamic nuclei (channels) |
|---|---|---|---|---|---|---|---|---|---|---|
| Thalamus* | 5 | 9 | 8 | 11 | 7 | 6 | 6 | 46 | 0 | Medial pulvinar (PuM)* (21); Ventral posterolateral (VPL)* (8); Lateral pulvinar (PuL)* (7); Centromedian (CM) (4); Ventral lateral (VL) (2); Anterior pulvinar (PuA) (1); Central lateral (CL) (1); Ventral anterior (VA) (1); Anterior nucleus (ANT) (1). Families: Pulvinar* (PuM, PuL, PuA; 29); Intralaminar* (CM, CL; 5); Lateral* (VA, VL, VPL; 11) |
| White matter (control)* | 9 | 12 | 9 | 19 | 27 | 30 | 6 | 106 | 0 | — |
| Spanning two systems or unlabeled | 10 | 8 | 3 | 6 | 12 | 4 | 6 | 43 | 6 | — |
| All intracranial channels | 150 | 149 | 97 | 124 | 143 | 128 | 6 | 791 | 78 | — |

**Supplementary Table S3. Paired comparisons of the two classifiers over the whole recording.** Each cell gives the participant-level mean difference (RF + HMM minus SleepSEEG) with bootstrap 95% CI, the number of participants with a positive difference and the Benjamini–Hochberg-corrected q of the recording-level mixed model. On the full intracranial montage, the two classifiers agreed equally well with the expert, but RF + HMM classified wake better on thalamic and white-matter channels and separated REM from the other stages better on every unit ($q \leq 0.040$ except temporal neocortex, $q = 0.082$), most markedly on the thalamus, where the REM AUC of SleepSEEG was 0.49. At night, no difference in κ, wake F1 or REM AUC was significant after correction ($q \geq 0.29$).

| Unit | Five-stage κ | Wake F1 | REM AUC |
|---|---|---|---|
| All intracranial channels | −0.014 [−0.092, +0.063]; 4/6, q = 0.664 | −0.002 [−0.073, +0.070]; 4/6, q = 0.931 | +0.117 [+0.037, +0.206]; 5/6, q = 0.032 |
| Mesial temporal | −0.046 [−0.120, +0.020]; 3/6, q = 0.568 | +0.013 [−0.043, +0.075]; 3/6, q = 0.679 | +0.106 [+0.026, +0.198]; 5/6, q = 0.040 |
| Temporal neocortex | +0.021 [−0.053, +0.087]; 4/6, q = 0.664 | +0.032 [−0.024, +0.097]; 5/6, q = 0.371 | +0.075 [+0.016, +0.154]; 5/6, q = 0.082 |
| Thalamus | +0.105 [+0.031, +0.171]; 5/6, q = 0.058 | +0.134 [+0.057, +0.205]; 5/6, q = 0.006 | +0.372 [+0.244, +0.487]; 6/6, q < 0.001 |
| White matter | +0.168 [+0.099, +0.249]; 6/6, q = 0.003 | +0.141 [+0.080, +0.203]; 6/6, q = 0.004 | +0.147 [+0.089, +0.213]; 6/6, q = 0.002 |

**Supplementary Table S4. N2+N3 detection and five-stage agreement by unit and classifier, whole recording.** Values are means across participants with bootstrap 95% CIs (2,000 resamples of participants); n, number of participants. For each system, “All channels” pools every channel of the system and the following rows are its structures or nuclei sampled in at least three participants. SleepSEEG is not defined for scalp or for scalp + thalamus fusion.

| System | Unit | n | Classifier | N2+N3 AUC | N2+N3 F1 | N2+N3 precision | N2+N3 sensitivity | Five-stage κ |
|---|---|---|---|---|---|---|---|---|
| Scalp | Scalp | 6 | RF + HMM | 0.96 [0.94, 0.97] | 0.72 [0.61, 0.82] | 0.78 [0.65, 0.88] | 0.75 [0.62, 0.86] | 0.59 [0.49, 0.70] |
| All intracranial | All channels | 6 | RF + HMM | 0.91 [0.84, 0.96] | 0.72 [0.60, 0.83] | 0.79 [0.64, 0.89] | 0.70 [0.60, 0.81] | 0.57 [0.45, 0.70] |
| | | | SleepSEEG | 0.90 [0.85, 0.95] | 0.78 [0.65, 0.87] | 0.81 [0.64, 0.92] | 0.78 [0.70, 0.83] | 0.58 [0.45, 0.69] |
| Mesial temporal lobe | All channels | 6 | RF + HMM | 0.88 [0.81, 0.94] | 0.62 [0.52, 0.74] | 0.71 [0.56, 0.82] | 0.64 [0.51, 0.78] | 0.47 [0.39, 0.56] |
| | | | SleepSEEG | 0.88 [0.83, 0.93] | 0.73 [0.60, 0.83] | 0.75 [0.60, 0.87] | 0.73 [0.62, 0.82] | 0.51 [0.41, 0.61] |
| | Hippocampus | 6 | RF + HMM | 0.87 [0.81, 0.93] | 0.61 [0.50, 0.72] | 0.69 [0.56, 0.79] | 0.63 [0.50, 0.79] | 0.46 [0.38, 0.55] |
| | | | SleepSEEG | 0.87 [0.80, 0.93] | 0.69 [0.57, 0.80] | 0.73 [0.58, 0.86] | 0.68 [0.58, 0.79] | 0.45 [0.34, 0.56] |
| | Amygdala | 4 | RF + HMM | 0.89 [0.82, 0.95] | 0.66 [0.53, 0.77] | 0.81 [0.79, 0.84] | 0.62 [0.48, 0.77] | 0.48 [0.37, 0.58] |
| | | | SleepSEEG | 0.87 [0.79, 0.95] | 0.75 [0.70, 0.81] | 0.80 [0.72, 0.88] | 0.73 [0.65, 0.80] | 0.45 [0.34, 0.58] |
| | Parahippocampal gyrus | 4 | RF + HMM | 0.81 [0.75, 0.86] | 0.52 [0.45, 0.59] | 0.62 [0.43, 0.82] | 0.52 [0.44, 0.62] | 0.36 [0.31, 0.43] |
| | | | SleepSEEG | 0.84 [0.77, 0.92] | 0.63 [0.49, 0.78] | 0.65 [0.47, 0.83] | 0.64 [0.51, 0.74] | 0.34 [0.24, 0.44] |
| Temporal neocortex | All channels | 6 | RF + HMM | 0.88 [0.80, 0.94] | 0.65 [0.54, 0.74] | 0.80 [0.64, 0.91] | 0.59 [0.50, 0.67] | 0.50 [0.42, 0.58] |
| | | | SleepSEEG | 0.85 [0.80, 0.90] | 0.70 [0.59, 0.77] | 0.76 [0.61, 0.88] | 0.68 [0.63, 0.72] | 0.48 [0.37, 0.57] |
| | Middle temporal gyrus | 6 | RF + HMM | 0.87 [0.79, 0.93] | 0.63 [0.53, 0.72] | 0.75 [0.59, 0.86] | 0.58 [0.50, 0.66] | 0.48 [0.40, 0.56] |

| System | Unit | n | Classifier | N2+N3 AUC | N2+N3 F1 | N2+N3 precision | N2+N3 sensitivity | Five-stage κ |
|---|---|---|---|---|---|---|---|---|
| | | | SleepSEEG | 0.81 [0.76, 0.87] | 0.66 [0.56, 0.73] | 0.68 [0.56, 0.78] | 0.67 [0.61, 0.72] | 0.42 [0.32, 0.51] |
| | Superior temporal gyrus | 6 | RF + HMM | 0.87 [0.80, 0.93] | 0.63 [0.49, 0.75] | 0.71 [0.57, 0.83] | 0.61 [0.48, 0.75] | 0.44 [0.36, 0.54] |
| | | | SleepSEEG | 0.80 [0.66, 0.92] | 0.62 [0.46, 0.77] | 0.64 [0.45, 0.83] | 0.64 [0.53, 0.75] | 0.30 [0.16, 0.42] |
| | Fusiform gyrus | 5 | RF + HMM | 0.87 [0.81, 0.93] | 0.64 [0.53, 0.76] | 0.72 [0.53, 0.86] | 0.63 [0.54, 0.73] | 0.49 [0.38, 0.60] |
| | | | SleepSEEG | 0.85 [0.80, 0.91] | 0.65 [0.53, 0.75] | 0.73 [0.55, 0.88] | 0.61 [0.54, 0.67] | 0.38 [0.22, 0.52] |
| | Inferior temporal gyrus | 4 | RF + HMM | 0.80 [0.68, 0.90] | 0.57 [0.37, 0.72] | 0.62 [0.40, 0.81] | 0.54 [0.38, 0.68] | 0.47 [0.33, 0.58] |
| | | | SleepSEEG | 0.80 [0.75, 0.86] | 0.57 [0.46, 0.67] | 0.61 [0.44, 0.76] | 0.57 [0.51, 0.61] | 0.34 [0.20, 0.48] |
| | Temporal pole, middle part | 3 | RF + HMM | 0.85 [0.81, 0.92] | 0.57 [0.49, 0.69] | 0.76 [0.58, 0.88] | 0.49 [0.43, 0.58] | 0.42 [0.31, 0.55] |
| | | | SleepSEEG | 0.78 [0.75, 0.81] | 0.61 [0.52, 0.66] | 0.70 [0.50, 0.83] | 0.56 [0.55, 0.59] | 0.34 [0.23, 0.47] |
| Frontal | All channels | 5 | RF + HMM | 0.93 [0.89, 0.96] | 0.75 [0.67, 0.82] | 0.86 [0.79, 0.90] | 0.72 [0.61, 0.82] | 0.56 [0.42, 0.69] |
| | | | SleepSEEG | 0.89 [0.79, 0.95] | 0.79 [0.65, 0.87] | 0.80 [0.63, 0.92] | 0.80 [0.71, 0.86] | 0.50 [0.37, 0.63] |
| | Gyrus rectus | 4 | RF + HMM | 0.92 [0.88, 0.96] | 0.75 [0.69, 0.80] | 0.82 [0.77, 0.88] | 0.75 [0.66, 0.85] | 0.53 [0.34, 0.68] |
| | | | SleepSEEG | 0.87 [0.79, 0.95] | 0.76 [0.69, 0.83] | 0.76 [0.68, 0.85] | 0.78 [0.70, 0.84] | 0.34 [0.22, 0.47] |
| | Inferior frontal gyrus, triangular part | 3 | RF + HMM | 0.93 [0.91, 0.94] | 0.78 [0.75, 0.80] | 0.80 [0.73, 0.86] | 0.78 [0.76, 0.80] | 0.64 [0.62, 0.68] |
| | | | SleepSEEG | 0.92 [0.87, 0.96] | 0.82 [0.76, 0.86] | 0.82 [0.74, 0.89] | 0.83 [0.78, 0.88] | 0.52 [0.49, 0.57] |
| | Medial orbitofrontal cortex | 3 | RF + HMM | 0.92 [0.84, 0.97] | 0.73 [0.50, 0.85] | 0.85 [0.82, 0.87] | 0.73 [0.52, 0.86] | 0.52 [0.26, 0.69] |

| System | Unit | n | Classifier | N2+N3 AUC | N2+N3 F1 | N2+N3 precision | N2+N3 sensitivity | Five-stage κ |
|---|---|---|---|---|---|---|---|---|
| | | | SleepSEEG | 0.94 [0.91, 0.97] | 0.83 [0.76, 0.88] | 0.85 [0.76, 0.92] | 0.83 [0.79, 0.90] | 0.46 [0.36, 0.52] |
| Insula / operculum | All channels | 4 | RF + HMM | 0.92 [0.83, 0.97] | 0.72 [0.53, 0.83] | 0.69 [0.47, 0.82] | 0.81 [0.71, 0.89] | 0.57 [0.40, 0.69] |
| | | | SleepSEEG | 0.88 [0.72, 0.97] | 0.72 [0.47, 0.88] | 0.74 [0.49, 0.91] | 0.73 [0.53, 0.88] | 0.31 [0.19, 0.44] |
| | Insula | 4 | RF + HMM | 0.92 [0.83, 0.98] | 0.72 [0.53, 0.83] | 0.69 [0.47, 0.83] | 0.81 [0.71, 0.87] | 0.55 [0.40, 0.63] |
| | | | SleepSEEG | 0.88 [0.72, 0.96] | 0.71 [0.47, 0.84] | 0.72 [0.46, 0.89] | 0.74 [0.53, 0.88] | 0.32 [0.19, 0.44] |
| Cingulate | All channels | 3 | RF + HMM | 0.96 [0.93, 0.99] | 0.82 [0.77, 0.87] | 0.85 [0.82, 0.87] | 0.82 [0.71, 0.88] | 0.60 [0.47, 0.79] |
| | | | SleepSEEG | 0.94 [0.89, 0.97] | 0.83 [0.77, 0.88] | 0.88 [0.79, 0.93] | 0.79 [0.76, 0.83] | 0.33 [0.22, 0.52] |
| Occipital | All channels | 3 | RF + HMM | 0.88 [0.75, 0.98] | 0.68 [0.48, 0.86] | 0.72 [0.53, 0.82] | 0.73 [0.60, 0.90] | 0.52 [0.40, 0.72] |
| | | | SleepSEEG | 0.89 [0.84, 0.95] | 0.70 [0.47, 0.85] | 0.71 [0.44, 0.89] | 0.71 [0.53, 0.83] | 0.35 [0.21, 0.50] |
| | Lingual gyrus | 3 | RF + HMM | 0.87 [0.72, 0.98] | 0.68 [0.45, 0.86] | 0.65 [0.37, 0.82] | 0.75 [0.57, 0.90] | 0.49 [0.34, 0.72] |
| | | | SleepSEEG | 0.86 [0.77, 0.95] | 0.66 [0.44, 0.85] | 0.68 [0.43, 0.89] | 0.66 [0.50, 0.83] | 0.27 [0.19, 0.36] |
| Thalamus | All channels | 6 | RF + HMM | 0.94 [0.90, 0.98] | 0.74 [0.62, 0.84] | 0.76 [0.63, 0.86] | 0.78 [0.66, 0.91] | 0.52 [0.41, 0.63] |
| | | | SleepSEEG | 0.93 [0.89, 0.96] | 0.78 [0.69, 0.87] | 0.79 [0.67, 0.89] | 0.81 [0.74, 0.87] | 0.42 [0.33, 0.49] |
| | Pulvinar family | 6 | RF + HMM | 0.93 [0.87, 0.97] | 0.73 [0.61, 0.83] | 0.74 [0.59, 0.85] | 0.79 [0.66, 0.91] | 0.49 [0.40, 0.58] |
| | | | SleepSEEG | 0.91 [0.87, 0.95] | 0.75 [0.65, 0.85] | 0.76 [0.63, 0.87] | 0.78 [0.70, 0.87] | 0.38 [0.28, 0.47] |
| | Medial pulvinar | 5 | RF + HMM | 0.92 [0.85, 0.98] | 0.71 [0.56, 0.86] | 0.74 [0.58, 0.85] | 0.77 [0.60, 0.93] | 0.47 [0.36, 0.58] |

| System | Unit | n | Classifier | N2+N3 AUC | N2+N3 F1 | N2+N3 precision | N2+N3 sensitivity | Five-stage κ |
|---|---|---|---|---|---|---|---|---|
| | | | SleepSEEG | 0.90 [0.85, 0.95] | 0.76 [0.64, 0.87] | 0.76 [0.62, 0.88] | 0.79 [0.69, 0.90] | 0.39 [0.30, 0.46] |
| | Lateral pulvinar | 5 | RF + HMM | 0.91 [0.83, 0.97] | 0.73 [0.61, 0.83] | 0.72 [0.56, 0.85] | 0.79 [0.66, 0.92] | 0.51 [0.38, 0.63] |
| | | | SleepSEEG | 0.86 [0.79, 0.94] | 0.68 [0.51, 0.82] | 0.68 [0.44, 0.86] | 0.74 [0.65, 0.82] | 0.25 [0.15, 0.35] |
| | Ventral posterolateral | 3 | RF + HMM | 0.94 [0.93, 0.96] | 0.71 [0.62, 0.76] | 0.68 [0.48, 0.81] | 0.85 [0.75, 0.97] | 0.50 [0.38, 0.63] |
| | | | SleepSEEG | 0.92 [0.90, 0.95] | 0.68 [0.58, 0.79] | 0.67 [0.55, 0.81] | 0.74 [0.67, 0.80] | 0.28 [0.19, 0.38] |
| | Intralaminar family | 3 | RF + HMM | 0.88 [0.79, 0.94] | 0.68 [0.57, 0.75] | 0.70 [0.49, 0.89] | 0.71 [0.66, 0.79] | 0.41 [0.27, 0.53] |
| | | | SleepSEEG | 0.86 [0.82, 0.94] | 0.62 [0.50, 0.76] | 0.61 [0.46, 0.76] | 0.68 [0.61, 0.80] | 0.20 [0.13, 0.25] |
| | Lateral thalamus family | 4 | RF + HMM | 0.95 [0.93, 0.97] | 0.75 [0.65, 0.85] | 0.72 [0.57, 0.82] | 0.86 [0.75, 0.96] | 0.54 [0.41, 0.67] |
| | | | SleepSEEG | 0.92 [0.89, 0.94] | 0.70 [0.62, 0.79] | 0.74 [0.59, 0.89] | 0.72 [0.67, 0.77] | 0.28 [0.21, 0.35] |
| White matter (control) | All channels | 6 | RF + HMM | 0.90 [0.83, 0.96] | 0.72 [0.62, 0.82] | 0.79 [0.64, 0.89] | 0.71 [0.61, 0.81] | 0.56 [0.45, 0.69] |
| | | | SleepSEEG | 0.87 [0.81, 0.92] | 0.69 [0.58, 0.79] | 0.70 [0.55, 0.84] | 0.73 [0.63, 0.80] | 0.40 [0.31, 0.47] |
| Fusion | Scalp + thalamus | 6 | RF + HMM | 0.97 [0.94, 0.98] | 0.73 [0.60, 0.84] | 0.81 [0.70, 0.89] | 0.76 [0.62, 0.90] | 0.62 [0.50, 0.75] |

**Supplementary Table S5. N2+N3 detection and five-stage agreement by unit and classifier, night.** Values are means across participants with bootstrap 95% CIs (2,000 resamples of participants); n, number of participants. For each system, "All channels" pools every channel of the system and the following rows are its structures or nuclei sampled in at least three participants. SleepSEEG is not defined for scalp or for scalp + thalamus fusion.

| System | Unit | n | Classifier | N2+N3 AUC | N2+N3 F1 | N2+N3 precision | N2+N3 sensitivity | Five-stage κ |
|---|---|---|---|---|---|---|---|---|
| Scalp | Scalp | 6 | RF + HMM | 0.88 [0.81, 0.94] | 0.74 [0.62, 0.85] | 0.82 [0.70, 0.91] | 0.75 [0.62, 0.87] | 0.43 [0.31, 0.56] |
| All intracranial | All channels | 6 | RF + HMM | 0.85 [0.76, 0.92] | 0.73 [0.64, 0.84] | 0.84 [0.76, 0.90] | 0.71 [0.61, 0.82] | 0.40 [0.27, 0.56] |
| | | | SleepSEEG | 0.82 [0.67, 0.92] | 0.80 [0.66, 0.88] | 0.83 [0.68, 0.94] | 0.79 [0.70, 0.84] | 0.44 [0.28, 0.58] |
| Mesial temporal lobe | All channels | 6 | RF + HMM | 0.82 [0.72, 0.91] | 0.64 [0.51, 0.77] | 0.81 [0.71, 0.87] | 0.63 [0.49, 0.78] | 0.32 [0.22, 0.44] |
| | | | SleepSEEG | 0.81 [0.66, 0.91] | 0.76 [0.62, 0.87] | 0.81 [0.66, 0.92] | 0.74 [0.62, 0.83] | 0.41 [0.26, 0.54] |
| | Hippocampus | 6 | RF + HMM | 0.81 [0.72, 0.89] | 0.64 [0.50, 0.77] | 0.80 [0.70, 0.87] | 0.62 [0.47, 0.77] | 0.30 [0.21, 0.40] |
| | | | SleepSEEG | 0.80 [0.66, 0.91] | 0.73 [0.60, 0.84] | 0.80 [0.64, 0.91] | 0.69 [0.58, 0.80] | 0.36 [0.23, 0.50] |
| | Amygdala | 4 | RF + HMM | 0.85 [0.77, 0.93] | 0.68 [0.51, 0.80] | 0.89 [0.86, 0.94] | 0.61 [0.45, 0.76] | 0.35 [0.19, 0.54] |
| | | | SleepSEEG | 0.85 [0.76, 0.94] | 0.80 [0.75, 0.85] | 0.90 [0.87, 0.94] | 0.73 [0.67, 0.80] | 0.43 [0.30, 0.55] |
| | Parahippocampal gyrus | 4 | RF + HMM | 0.76 [0.61, 0.90] | 0.57 [0.47, 0.69] | 0.83 [0.65, 0.94] | 0.51 [0.45, 0.61] | 0.28 [0.16, 0.45] |
| | | | SleepSEEG | 0.78 [0.60, 0.92] | 0.70 [0.53, 0.84] | 0.78 [0.56, 0.92] | 0.65 [0.51, 0.77] | 0.31 [0.17, 0.45] |
| Temporal neocortex | All channels | 6 | RF + HMM | 0.81 [0.72, 0.89] | 0.65 [0.53, 0.74] | 0.87 [0.79, 0.93] | 0.57 [0.49, 0.66] | 0.31 [0.22, 0.41] |
| | | | SleepSEEG | 0.78 [0.64, 0.86] | 0.73 [0.62, 0.79] | 0.82 [0.67, 0.92] | 0.69 [0.62, 0.73] | 0.36 [0.23, 0.46] |
| | Middle temporal gyrus | 6 | RF + HMM | 0.80 [0.71, 0.87] | 0.66 [0.55, 0.73] | 0.86 [0.76, 0.93] | 0.57 [0.49, 0.65] | 0.31 [0.22, 0.40] |

| System | Unit | n | Classifier | N2+N3 AUC | N2+N3 F1 | N2+N3 precision | N2+N3 sensitivity | Five-stage κ |
|---|---|---|---|---|---|---|---|---|
| | | | SleepSEEG | 0.75 [0.61, 0.85] | 0.70 [0.59, 0.78] | 0.77 [0.63, 0.88] | 0.67 [0.60, 0.72] | 0.32 [0.21, 0.41] |
| | Superior temporal gyrus | 6 | RF + HMM | 0.81 [0.72, 0.90] | 0.65 [0.51, 0.78] | 0.83 [0.77, 0.89] | 0.61 [0.46, 0.75] | 0.31 [0.21, 0.42] |
| | | | SleepSEEG | 0.74 [0.58, 0.88] | 0.68 [0.55, 0.82] | 0.74 [0.58, 0.88] | 0.65 [0.52, 0.77] | 0.27 [0.11, 0.42] |
| | Fusiform gyrus | 5 | RF + HMM | 0.83 [0.74, 0.88] | 0.70 [0.61, 0.78] | 0.82 [0.63, 0.94] | 0.65 [0.57, 0.74] | 0.36 [0.22, 0.48] |
| | | | SleepSEEG | 0.81 [0.72, 0.89] | 0.71 [0.59, 0.78] | 0.83 [0.67, 0.92] | 0.63 [0.56, 0.69] | 0.34 [0.21, 0.46] |
| | Inferior temporal gyrus | 4 | RF + HMM | 0.73 [0.54, 0.87] | 0.63 [0.42, 0.76] | 0.82 [0.64, 0.94] | 0.53 [0.36, 0.66] | 0.35 [0.15, 0.54] |
| | | | SleepSEEG | 0.70 [0.54, 0.79] | 0.63 [0.49, 0.71] | 0.73 [0.54, 0.87] | 0.56 [0.48, 0.62] | 0.26 [0.13, 0.35] |
| | Temporal pole, middle part | 3 | RF + HMM | 0.81 [0.74, 0.87] | 0.62 [0.53, 0.69] | 0.93 [0.90, 0.99] | 0.48 [0.41, 0.56] | 0.32 [0.20, 0.49] |
| | | | SleepSEEG | 0.75 [0.73, 0.78] | 0.67 [0.66, 0.69] | 0.87 [0.87, 0.87] | 0.55 [0.54, 0.58] | 0.30 [0.27, 0.32] |
| Frontal | All channels | 5 | RF + HMM | 0.89 [0.84, 0.94] | 0.76 [0.66, 0.85] | 0.90 [0.87, 0.92] | 0.72 [0.61, 0.82] | 0.41 [0.26, 0.60] |
| | | | SleepSEEG | 0.87 [0.78, 0.94] | 0.84 [0.76, 0.90] | 0.88 [0.79, 0.94] | 0.81 [0.74, 0.88] | 0.46 [0.34, 0.60] |
| | Gyrus rectus | 4 | RF + HMM | 0.89 [0.85, 0.93] | 0.79 [0.73, 0.86] | 0.89 [0.85, 0.93] | 0.75 [0.66, 0.87] | 0.38 [0.21, 0.60] |
| | | | SleepSEEG | 0.85 [0.74, 0.94] | 0.81 [0.73, 0.88] | 0.86 [0.80, 0.92] | 0.78 [0.69, 0.86] | 0.38 [0.26, 0.50] |
| | Inferior frontal gyrus, triangular part | 3 | RF + HMM | 0.91 [0.84, 0.96] | 0.84 [0.80, 0.89] | 0.90 [0.84, 0.93] | 0.80 [0.76, 0.86] | 0.54 [0.44, 0.72] |
| | | | SleepSEEG | 0.91 [0.84, 0.96] | 0.87 [0.81, 0.91] | 0.91 [0.87, 0.94] | 0.84 [0.76, 0.89] | 0.52 [0.39, 0.68] |
| | Medial orbitofrontal cortex | 3 | RF + HMM | 0.91 [0.83, 0.95] | 0.75 [0.49, 0.88] | 0.89 [0.88, 0.90] | 0.73 [0.48, 0.86] | 0.38 [0.15, 0.60] |

| System | Unit | n | Classifier | N2+N3 AUC | N2+N3 F1 | N2+N3 precision | N2+N3 sensitivity | Five-stage κ |
|---|---|---|---|---|---|---|---|---|
| | | | SleepSEEG | 0.93 [0.91, 0.96] | 0.88 [0.86, 0.92] | 0.93 [0.90, 0.94] | 0.85 [0.80, 0.91] | 0.47 [0.40, 0.57] |
| Insula / operculum | All channels | 4 | RF + HMM | 0.83 [0.67, 0.94] | 0.76 [0.59, 0.87] | 0.75 [0.55, 0.87] | 0.83 [0.75, 0.91] | 0.38 [0.14, 0.61] |
| | | | SleepSEEG | 0.84 [0.65, 0.95] | 0.76 [0.54, 0.90] | 0.81 [0.57, 0.95] | 0.74 [0.54, 0.88] | 0.36 [0.14, 0.52] |
| | Insula | 4 | RF + HMM | 0.83 [0.67, 0.94] | 0.76 [0.59, 0.87] | 0.76 [0.56, 0.88] | 0.82 [0.75, 0.89] | 0.39 [0.14, 0.61] |
| | | | SleepSEEG | 0.83 [0.65, 0.94] | 0.75 [0.54, 0.87] | 0.79 [0.56, 0.94] | 0.73 [0.54, 0.86] | 0.35 [0.14, 0.51] |
| Cingulate | All channels | 3 | RF + HMM | 0.92 [0.84, 0.97] | 0.84 [0.75, 0.89] | 0.87 [0.85, 0.91] | 0.82 [0.69, 0.90] | 0.45 [0.32, 0.69] |
| | | | SleepSEEG | 0.92 [0.82, 0.97] | 0.85 [0.79, 0.92] | 0.90 [0.79, 0.96] | 0.82 [0.78, 0.88] | 0.40 [0.35, 0.45] |
| Occipital | All channels | 3 | RF + HMM | 0.81 [0.68, 0.93] | 0.70 [0.52, 0.88] | 0.76 [0.62, 0.86] | 0.72 [0.60, 0.91] | 0.31 [0.17, 0.52] |
| | | | SleepSEEG | 0.78 [0.55, 0.94] | 0.72 [0.48, 0.88] | 0.75 [0.47, 0.94] | 0.71 [0.52, 0.84] | 0.28 [0.12, 0.40] |
| | Lingual gyrus | 3 | RF + HMM | 0.81 [0.67, 0.93] | 0.68 [0.44, 0.88] | 0.74 [0.56, 0.86] | 0.70 [0.49, 0.91] | 0.26 [0.10, 0.52] |
| | | | SleepSEEG | 0.75 [0.53, 0.94] | 0.69 [0.46, 0.88] | 0.74 [0.47, 0.94] | 0.67 [0.49, 0.84] | 0.25 [0.11, 0.40] |
| Thalamus | All channels | 6 | RF + HMM | 0.88 [0.81, 0.94] | 0.76 [0.64, 0.86] | 0.78 [0.64, 0.88] | 0.79 [0.67, 0.91] | 0.31 [0.21, 0.42] |
| | | | SleepSEEG | 0.86 [0.80, 0.93] | 0.80 [0.71, 0.88] | 0.82 [0.70, 0.90] | 0.82 [0.75, 0.88] | 0.38 [0.26, 0.50] |
| | Pulvinar family | 6 | RF + HMM | 0.87 [0.79, 0.94] | 0.75 [0.64, 0.85] | 0.77 [0.64, 0.87] | 0.80 [0.67, 0.92] | 0.28 [0.19, 0.38] |
| | | | SleepSEEG | 0.83 [0.76, 0.91] | 0.78 [0.68, 0.87] | 0.80 [0.68, 0.89] | 0.79 [0.70, 0.87] | 0.33 [0.21, 0.45] |
| | Medial pulvinar | 5 | RF + HMM | 0.86 [0.78, 0.95] | 0.73 [0.58, 0.88] | 0.77 [0.62, 0.88] | 0.78 [0.61, 0.94] | 0.29 [0.17, 0.42] |

| System | Unit | n | Classifier | N2+N3 AUC | N2+N3 F1 | N2+N3 precision | N2+N3 sensitivity | Five-stage κ |
|---|---|---|---|---|---|---|---|---|
| | | | SleepSEEG | 0.85 [0.76, 0.93] | 0.79 [0.69, 0.88] | 0.80 [0.67, 0.90] | 0.80 [0.70, 0.90] | 0.33 [0.22, 0.43] |
| | Lateral pulvinar | 5 | RF + HMM | 0.86 [0.76, 0.94] | 0.76 [0.64, 0.86] | 0.76 [0.60, 0.86] | 0.81 [0.69, 0.93] | 0.33 [0.22, 0.46] |
| | | | SleepSEEG | 0.81 [0.70, 0.92] | 0.75 [0.62, 0.87] | 0.77 [0.61, 0.89] | 0.75 [0.67, 0.84] | 0.27 [0.16, 0.38] |
| | Ventral posterolateral | 3 | RF + HMM | 0.84 [0.80, 0.89] | 0.74 [0.65, 0.80] | 0.71 [0.51, 0.82] | 0.86 [0.73, 1.00] | 0.25 [0.09, 0.39] |
| | | | SleepSEEG | 0.82 [0.77, 0.89] | 0.72 [0.60, 0.82] | 0.74 [0.59, 0.85] | 0.74 [0.64, 0.81] | 0.22 [0.09, 0.34] |
| | Intralaminar family | 3 | RF + HMM | 0.81 [0.71, 0.88] | 0.71 [0.59, 0.79] | 0.73 [0.50, 0.89] | 0.73 [0.65, 0.82] | 0.24 [0.12, 0.35] |
| | | | SleepSEEG | 0.79 [0.73, 0.86] | 0.69 [0.58, 0.80] | 0.73 [0.58, 0.85] | 0.69 [0.63, 0.79] | 0.21 [0.11, 0.31] |
| | Lateral thalamus family | 4 | RF + HMM | 0.87 [0.81, 0.93] | 0.77 [0.68, 0.85] | 0.74 [0.58, 0.83] | 0.88 [0.78, 0.97] | 0.35 [0.15, 0.58] |
| | | | SleepSEEG | 0.85 [0.78, 0.91] | 0.75 [0.65, 0.83] | 0.79 [0.66, 0.94] | 0.74 [0.67, 0.79] | 0.28 [0.15, 0.39] |
| White matter (control) | All channels | 6 | RF + HMM | 0.84 [0.74, 0.92] | 0.75 [0.65, 0.84] | 0.84 [0.76, 0.89] | 0.72 [0.64, 0.83] | 0.41 [0.27, 0.57] |
| | | | SleepSEEG | 0.80 [0.66, 0.91] | 0.76 [0.63, 0.85] | 0.80 [0.64, 0.91] | 0.74 [0.64, 0.82] | 0.38 [0.24, 0.50] |
| Fusion | Scalp + thalamus | 6 | RF + HMM | 0.89 [0.82, 0.95] | 0.73 [0.57, 0.87] | 0.83 [0.74, 0.89] | 0.76 [0.60, 0.90] | 0.46 [0.32, 0.61] |

**Supplementary Table S6. Thalamus versus other units: paired comparisons.** Each cell gives the participant-level mean difference (thalamus minus the unit in the row; last row, scalp + thalamus minus scalp), its bootstrap 95% CI, and the number of participants with a positive difference with the Benjamini–Hochberg-corrected q of the recording-level mixed model. With RF + HMM the thalamus was close to scalp in N2+N3 AUC and exceeded the mesial temporal lobe, hippocampus and temporal neocortex in all six participants over the whole recording (five or six of six at night); its five-stage κ was lower than that of scalp in all six participants and similar to that of the other intracranial units, and its N3 F1 was lower than that of every other unit. With SleepSEEG, thalamic N2+N3 AUC was higher than that of every unit but not significantly after correction (q ≥ 0.08), and F1 was significantly higher than that of the hippocampus, temporal neocortex and white matter. Adding the thalamus to scalp raised N2+N3 AUC in five of six participants. SleepSEEG is not defined for scalp.

| Classifier | Thalamus vs | N2+N3 AUC Whole | N2+N3 AUC Night | N2+N3 F1 Whole | N2+N3 F1 Night | Five-stage κ Whole | N3 F1 Whole |
|---|---|---|---|---|---|---|---|
| RF + HMM | Scalp | −0.015<br>[−0.037, +0.004]<br>3/6, q = 0.156 | −0.004<br>[−0.030, +0.022]<br>4/6, q = 0.665 | +0.017<br>[−0.010, +0.042]<br>4/6, q = 0.501 | +0.013<br>[−0.015, +0.046]<br>4/6, q = 0.649 | −0.073<br>[−0.113, −0.033]<br>0/6, q = 0.013 | −0.182<br>[−0.305, −0.059]<br>1/6, q = 0.004 |
| | All intracranial channels | +0.032<br>[+0.007, +0.067]<br>5/6, q = 0.064 | +0.030<br>[+0.001, +0.060]<br>4/6, q = 0.068 | +0.017<br>[+0.001, +0.033]<br>5/6, q = 0.286 | +0.021<br>[+0.001, +0.043]<br>4/6, q = 0.214 | −0.044<br>[−0.099, +0.016]<br>2/6, q = 0.401 | −0.209<br>[−0.317, −0.100]<br>1/6, q = 0.001 |
| | Mesial temporal lobe | +0.068<br>[+0.027, +0.111]<br>6/6, q = 0.024 | +0.061<br>[+0.022, +0.105]<br>6/6, q = 0.018 | +0.114<br>[+0.067, +0.169]<br>6/6, q = 0.008 | +0.111<br>[+0.062, +0.167]<br>6/6, q = 0.010 | +0.056<br>[−0.021, +0.138]<br>4/6, q = 0.468 | −0.154<br>[−0.249, −0.061]<br>1/6, q = 0.003 |
| | Hippocampus | +0.070<br>[+0.029, +0.114]<br>6/6, q = 0.024 | +0.072<br>[+0.029, +0.124]<br>6/6, q = 0.018 | +0.123<br>[+0.072, +0.181]<br>6/6, q = 0.004 | +0.120<br>[+0.071, +0.169]<br>6/6, q = 0.007 | +0.065<br>[−0.008, +0.152]<br>5/6, q = 0.401 | −0.175<br>[−0.271, −0.072]<br>1/6, q = 0.001 |
| | Temporal neocortex | +0.064<br>[+0.025, +0.107]<br>6/6, q = 0.024 | +0.066<br>[+0.021, +0.108]<br>5/6, q = 0.018 | +0.083<br>[+0.005, +0.148]<br>5/6, q = 0.060 | +0.106<br>[+0.021, +0.176]<br>5/6, q = 0.023 | +0.022<br>[−0.034, +0.085]<br>2/6, q = 0.736 | −0.209<br>[−0.323, −0.084]<br>1/6, q = 0.001 |
| | White matter (control) | +0.040<br>[+0.010, +0.075]<br>5/6, q = 0.036 | +0.035<br>[+0.003, +0.073]<br>5/6, q = 0.068 | +0.013<br>[−0.041, +0.055]<br>4/6, q = 0.501 | +0.009<br>[−0.040, +0.053]<br>3/6, q = 0.555 | −0.043<br>[−0.103, +0.026]<br>2/6, q = 0.401 | −0.195<br>[−0.315, −0.077]<br>1/6, q = 0.004 |
| SleepSEEG | All intracranial channels | +0.024<br>[−0.024, +0.077]<br>3/6, q = 0.400 | +0.042<br>[−0.027, +0.145]<br>4/6, q = 0.388 | +0.005<br>[−0.044, +0.053]<br>4/6, q = 0.934 | +0.009<br>[−0.035, +0.055]<br>4/6, q = 0.751 | −0.163<br>[−0.203, −0.123]<br>0/6, q < 0.001 | −0.081<br>[−0.131, −0.030]<br>2/6, q = 0.003 |
| | Mesial temporal lobe | +0.044<br>[−0.013, +0.104]<br>3/6, q = 0.243 | +0.058<br>[−0.015, +0.157]<br>5/6, q = 0.264 | +0.054<br>[+0.003, +0.104]<br>4/6, q = 0.105 | +0.040<br>[−0.011, +0.092]<br>5/6, q = 0.158 | −0.094<br>[−0.135, −0.067]<br>0/6, q = 0.002 | −0.068<br>[−0.153, +0.023]<br>2/6, q = 0.079 |

| Classifier | Thalamus vs | N2+N3 AUC Whole | N2+N3 AUC Night | N2+N3 F1 Whole | N2+N3 F1 Night | Five-stage κ Whole | N3 F1 Whole |
|---|---|---|---|---|---|---|---|
| | Hippocampus | +0.058 [−0.015, +0.142] 3/6, q = 0.243 | +0.068 [−0.015, +0.173] 5/6, q = 0.264 | +0.093 [+0.030, +0.158] 5/6, q = 0.039 | +0.071 [+0.024, +0.119] 6/6, q = 0.056 | −0.036 [−0.087, +0.029] 1/6, q = 0.191 | −0.052 [−0.121, +0.035] 2/6, q = 0.079 |
| | Temporal neocortex | +0.073 [+0.028, +0.127] 6/6, q = 0.084 | +0.088 [+0.022, +0.178] 5/6, q = 0.264 | +0.087 [+0.046, +0.125] 6/6, q = 0.004 | +0.074 [+0.027, +0.117] 5/6, q = 0.056 | −0.062 [−0.110, −0.020] 1/6, q = 0.049 | −0.064 [−0.114, −0.008] 2/6, q = 0.008 |
| | White matter (control) | +0.058 [+0.010, +0.104] 5/6, q = 0.084 | +0.060 [−0.007, +0.157] 5/6, q = 0.264 | +0.092 [+0.058, +0.123] 6/6, q = 0.002 | +0.049 [+0.016, +0.093] 5/6, q = 0.068 | +0.020 [−0.024, +0.060] 4/6, q = 0.661 | −0.063 [−0.107, −0.018] 1/6, q = 0.008 |
| RF + HMM | Scalp + thalamus vs scalp | +0.008 [+0.004, +0.011] 5/6, q = 0.011 | +0.009 [−0.003, +0.023] 4/6, q = 0.411 | +0.007 [−0.015, +0.032] 3/6, q = 0.963 | −0.014 [−0.050, +0.013] 3/6, q = 0.745 | +0.028 [+0.009, +0.048] 5/6, q = 0.121 | −0.020 [−0.083, +0.044] 2/6, q = 0.731 |

**Supplementary Table S7. Pretrained scalp scorers compared with RF + HMM on scalp.** Values are means across the six participants with bootstrap 95% CIs (recordings averaged within participant), on the same artifact- and seizure-free epochs as the other classifiers; F1 refers to the decoded labels. Δ, participant-level mean difference (scorer minus RF + HMM on scalp) with bootstrap 95% CI, the number of participants in whom the scorer was higher, and the Benjamini–Hochberg-corrected q of the recording-level mixed model, corrected across scorers within each window and metric.

| Window | Scorer | Five-stage κ | Δ κ | N2+N3 AUC | Δ AUC | N2+N3 F1 | Δ F1 |
|---|---|---|---|---|---|---|---|
| Whole recording | RF + HMM, scalp (reference) | 0.59 [0.49, 0.70] | — | 0.96 [0.94, 0.97] | — | 0.72 [0.61, 0.82] | — |
| | U-Sleep | 0.73 [0.67, 0.78] | +0.131 [+0.045, +0.249] 5/6, q = 0.039 | 0.96 [0.94, 0.98] | +0.007 [+0.003, +0.010] 5/6, q = 0.598 | 0.87 [0.79, 0.93] | +0.149 [+0.077, +0.258] 6/6, q = 0.009 |
| | DeepResNet | 0.73 [0.66, 0.79] | +0.135 [+0.073, +0.223] 6/6, q = 0.006 | 0.97 [0.95, 0.98] | +0.010 [+0.006, +0.015] 6/6, q = 0.391 | 0.87 [0.79, 0.92] | +0.146 [+0.076, +0.256] 6/6, q = 0.009 |
| | SleepTransformer | 0.75 [0.69, 0.80] | +0.155 [+0.097, +0.230] 6/6, q < 0.001 | 0.96 [0.95, 0.98] | +0.005 [−0.001, +0.012] 4/6, q = 0.668 | 0.86 [0.78, 0.92] | +0.142 [+0.070, +0.251] 6/6, q = 0.009 |
| | SLEEPYLAND ensemble | 0.76 [0.71, 0.82] | +0.170 [+0.107, +0.241] 6/6, q < 0.001 | 0.96 [0.94, 0.97] | −0.002 [−0.010, +0.007] 3/6, q = 0.978 | 0.88 [0.82, 0.93] | +0.161 [+0.083, +0.265] 6/6, q = 0.009 |
| Night | RF + HMM, scalp (reference) | 0.43 [0.30, 0.57] | — | 0.88 [0.82, 0.94] | — | 0.74 [0.63, 0.85] | — |
| | U-Sleep | 0.64 [0.53, 0.71] | +0.209 [+0.121, +0.299] 6/6, q < 0.001 | 0.92 [0.87, 0.96] | +0.035 [+0.016, +0.059] 6/6, q = 0.087 | 0.87 [0.79, 0.93] | +0.125 [+0.049, +0.245] 6/6, q = 0.040 |
| | DeepResNet | 0.63 [0.54, 0.72] | +0.208 [+0.127, +0.287] 6/6, q < 0.001 | 0.92 [0.87, 0.96] | +0.037 [+0.015, +0.062] 6/6, q = 0.087 | 0.87 [0.79, 0.93] | +0.124 [+0.050, +0.239] 6/6, q = 0.040 |
| | SleepTransformer | 0.64 [0.54, 0.72] | +0.210 [+0.134, +0.286] 6/6, q < 0.001 | 0.91 [0.86, 0.96] | +0.031 [+0.015, +0.051] 6/6, q = 0.087 | 0.87 [0.79, 0.93] | +0.127 [+0.050, +0.246] 6/6, q = 0.040 |
| | SLEEPYLAND ensemble | 0.66 [0.58, 0.73] | +0.238 [+0.159, +0.309] 6/6, q < 0.001 | 0.92 [0.87, 0.96] | +0.036 [+0.012, +0.064] 6/6, q = 0.087 | 0.88 [0.81, 0.93] | +0.138 [+0.057, +0.253] 6/6, q = 0.040 |

**Supplementary Table S8. Thalamus versus channel-count-matched subsets of other pools.** For each participant, up to 30 random subsets of other channels were drawn with as many channels as the participant's thalamic channels (5–11), averaged and scored. Each cell gives the participant-level mean difference (thalamus minus the matched subset), its bootstrap 95% CI, and the number of participants with a positive difference with the Benjamini–Hochberg-corrected q of the recording-level mixed model. Mesial temporal subsets were available in five participants, whose mesial temporal pool was at least as large as their thalamic count.

| **Classifier** | **Thalamus vs matched** | **N2+N3 AUC Whole** | **N2+N3 AUC Night** | **N2+N3 F1 Whole** | **N2+N3 F1 Night** |
|---|---|---|---|---|---|
| RF + HMM | All intracranial channels | +0.036 [+0.013, +0.063] 6/6, q = 0.011 | +0.036 [+0.007, +0.062] 5/6, q = 0.020 | +0.021 [−0.014, +0.050] 5/6, q = 0.272 | +0.029 [+0.001, +0.057] 5/6, q = 0.140 |
| | White matter | +0.043 [+0.015, +0.075] 5/6, q = 0.011 | +0.036 [+0.007, +0.069] 5/6, q = 0.030 | +0.017 [−0.042, +0.060] 5/6, q = 0.467 | +0.014 [−0.046, +0.058] 4/6, q = 0.497 |
| | Mesial temporal lobe | +0.082 [+0.035, +0.131] 5/5, q = 0.011 | +0.074 [+0.032, +0.118] 5/5, q = 0.011 | +0.127 [+0.066, +0.189] 5/5, q = 0.003 | +0.113 [+0.059, +0.164] 5/5, q = 0.016 |
| | Temporal neocortex | +0.069 [+0.033, +0.106] 6/6, q = 0.011 | +0.069 [+0.027, +0.112] 5/6, q = 0.011 | +0.084 [+0.003, +0.155] 5/6, q = 0.081 | +0.091 [+0.011, +0.155] 5/6, q = 0.047 |
| SleepSEEG | All intracranial channels | +0.047 [+0.003, +0.094] 4/6, q = 0.097 | +0.058 [−0.002, +0.143] 4/6, q = 0.176 | +0.054 [+0.004, +0.100] 4/6, q = 0.063 | +0.045 [+0.005, +0.085] 4/6, q = 0.069 |
| | White matter | +0.068 [+0.015, +0.113] 5/6, q = 0.029 | +0.069 [+0.002, +0.169] 5/6, q = 0.176 | +0.111 [+0.074, +0.146] 6/6, q < 0.001 | +0.061 [+0.025, +0.104] 5/6, q = 0.024 |
| | Mesial temporal lobe | +0.065 [−0.001, +0.130] 4/5, q = 0.105 | +0.072 [−0.016, +0.184] 4/5, q = 0.176 | +0.090 [+0.033, +0.146] 4/5, q = 0.020 | +0.062 [+0.003, +0.116] 4/5, q = 0.069 |
| | Temporal neocortex | +0.092 [+0.040, +0.146] 6/6, q = 0.018 | +0.100 [+0.033, +0.184] 5/6, q = 0.107 | +0.123 [+0.069, +0.176] 6/6, q < 0.001 | +0.096 [+0.045, +0.144] 6/6, q = 0.008 |

**Supplementary Table S9. Anatomical systems versus the white-matter control, whole recording.** Each cell gives the participant-level mean difference (system minus white matter), its bootstrap 95% CI, and the number of participants with a positive difference with the Benjamini–Hochberg-corrected q of the recording-level mixed model (comparison family: systems versus white matter, per classifier, window and metric); the number of participants sampled is given after each system. No system differed from white matter in N2+N3 AUC or F1 with either classifier. For five-stage κ, white matter matched or exceeded every gray-matter system with RF + HMM, whereas with SleepSEEG the mesial temporal lobe and temporal neocortex exceeded white matter in all six participants.

| Classifier | System (participants) | N2+N3 AUC | N2+N3 F1 | Five-stage κ |
|---|---|---|---|---|
| RF + HMM | Mesial temporal lobe (6) | −0.028<br>[−0.096, +0.025]<br>3/6, q = 0.511 | −0.101<br>[−0.185, −0.018]<br>2/6, q = 0.141 | −0.099<br>[−0.203, −0.026]<br>1/6, q = 0.099 |
| | Temporal neocortex (6) | −0.024<br>[−0.047, +0.009]<br>1/6, q = 0.268 | −0.070<br>[−0.132, −0.008]<br>1/6, q = 0.139 | −0.065<br>[−0.143, +0.018]<br>1/6, q = 0.188 |
| | Frontal (5) | −0.010<br>[−0.025, +0.007]<br>2/5, q = 0.288 | −0.018<br>[−0.059, +0.025]<br>1/5, q = 0.705 | −0.034<br>[−0.086, +0.022]<br>1/5, q = 0.329 |
| | Insula / operculum (4) | +0.020<br>[−0.010, +0.050]<br>2/4, q = 0.687 | −0.020<br>[−0.057, +0.036]<br>1/4, q = 0.705 | −0.063<br>[−0.112, +0.016]<br>1/4, q = 0.168 |
| | Cingulate (3) | +0.016<br>[−0.016, +0.054]<br>2/3, q = 0.488 | +0.016<br>[−0.031, +0.081]<br>1/3, q = 0.566 | −0.052<br>[−0.248, +0.130]<br>1/3, q = 0.604 |
| | Occipital (3) | +0.016<br>[−0.004, +0.044]<br>2/3, q = 0.288 | +0.008<br>[−0.022, +0.029]<br>2/3, q = 0.705 | −0.005<br>[−0.038, +0.026]<br>1/3, q = 0.932 |
| SleepSEEG | Mesial temporal lobe (6) | +0.014<br>[−0.027, +0.068]<br>3/6, q = 0.775 | +0.039<br>[+0.002, +0.084]<br>5/6, q = 0.253 | +0.114<br>[+0.062, +0.153]<br>6/6, q = 0.003 |
| | Temporal neocortex (6) | −0.015<br>[−0.052, +0.036]<br>1/6, q = 0.826 | +0.005<br>[−0.032, +0.041]<br>3/6, q = 0.858 | +0.082<br>[+0.045, +0.119]<br>6/6, q = 0.005 |
| | Frontal (5) | +0.002<br>[−0.029, +0.026]<br>3/5, q = 0.941 | +0.045<br>[−0.020, +0.098]<br>4/5, q = 0.367 | +0.069<br>[−0.028, +0.151]<br>4/5, q = 0.361 |

| **Classifier** | **System (participants)** | **N2+N3 AUC** | **N2+N3 F1** | **Five-stage κ** |
|---|---|---|---|---|
| | Insula / operculum (4) | −0.006<br>[−0.104, +0.064]<br>3/4, q = 0.941 | +0.025<br>[−0.055, +0.105]<br>2/4, q = 0.684 | −0.109<br>[−0.183, −0.047]<br>0/4, q = 0.066 |
| | Cingulate (3) | +0.053<br>[−0.005, +0.124]<br>2/3, q = 0.382 | +0.080<br>[−0.017, +0.163]<br>2/3, q = 0.253 | −0.118<br>[−0.303, +0.179]<br>1/3, q = 0.572 |
| | Occipital (3) | +0.053<br>[−0.005, +0.146]<br>2/3, q = 0.399 | +0.071<br>[−0.002, +0.173]<br>2/3, q = 0.277 | −0.003<br>[−0.165, +0.160]<br>1/3, q = 0.968 |

**Supplementary Table S10. Seizure-onset-zone (SOZ) analyses, whole recording.** The first block gives the effect of removing SOZ channels from each unit that contained them (unit without minus with SOZ channels; q, Benjamini–Hochberg-corrected p of the recording-level mixed model), in the participants whose channels in that unit included SOZ channels: three for temporal neocortex, six for the other units. The second block compares the thalamus with mesial temporal channels outside the SOZ (thalamus minus non-SOZ mesial temporal channels; p, Wilcoxon signed-rank test on participant means). Each cell gives the participant-level mean difference, its bootstrap 95% CI, and the number of participants with a positive difference with q or p. No thalamic channel lay in the SOZ.

| Analysis | Classifier | Unit | N2+N3 AUC | N2+N3 F1 |
|---|---|---|---|---|
| SOZ channels removed | RF + HMM | All intracranial channels | +0.002<br>[−0.000, +0.006]<br>5/6, q = 0.440 | +0.006<br>[+0.001, +0.012]<br>5/6, q = 0.138 |
| | | Mesial temporal lobe | −0.006<br>[−0.014, +0.001]<br>2/6, q = 0.440 | +0.000<br>[−0.025, +0.031]<br>3/6, q = 0.851 |
| | | Hippocampus | +0.002<br>[−0.013, +0.026]<br>1/6, q = 0.907 | +0.026<br>[−0.017, +0.072]<br>3/6, q = 0.322 |
| | | Temporal neocortex | +0.000<br>[−0.000, +0.001]<br>2/3, q = 0.907 | +0.002<br>[−0.003, +0.011]<br>1/3, q = 0.510 |
| | SleepSEEG | All intracranial channels | −0.003<br>[−0.009, +0.001]<br>1/6, q = 0.606 | −0.005<br>[−0.012, +0.003]<br>2/6, q = 0.459 |
| | | Mesial temporal lobe | −0.030<br>[−0.081, +0.004]<br>2/6, q = 0.511 | −0.032<br>[−0.082, +0.004]<br>2/6, q = 0.432 |
| | | Hippocampus | −0.030<br>[−0.075, +0.006]<br>2/6, q = 0.511 | −0.028<br>[−0.065, +0.005]<br>2/6, q = 0.432 |
| | | Temporal neocortex | −0.002<br>[−0.006, +0.003]<br>1/3, q = 0.653 | +0.002<br>[−0.001, +0.009]<br>1/3, q = 0.459 |
| Thalamus vs non-SOZ mesial temporal channels | RF + HMM | Thalamus vs non-SOZ mesial temporal lobe | +0.074<br>[+0.035, +0.122]<br>6/6, p = 0.031 | +0.113<br>[+0.047, +0.190]<br>6/6, p = 0.031 |

| Analysis | Classifier | Unit | N2+N3 AUC | N2+N3 F1 |
|---|---|---|---|---|
| | SleepSEEG | Thalamus vs non-SOZ mesial temporal lobe | +0.073<br>[+0.030, +0.117]<br>5/6, p = 0.062 | +0.086<br>[+0.051, +0.121]<br>6/6, p = 0.031 |

### Supplementary Figures

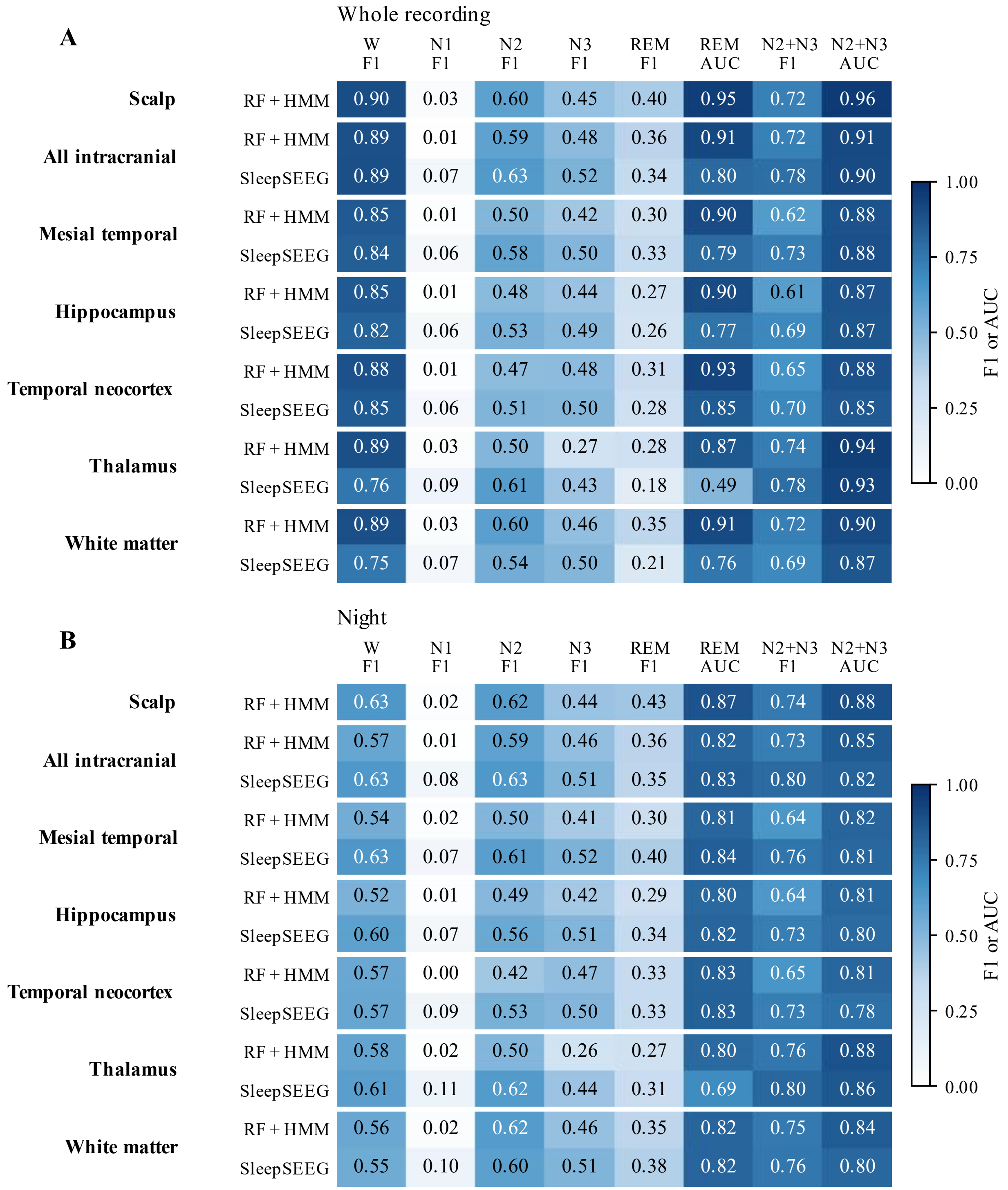


**Supplementary Figure S1. Per-stage detection by unit and classifier.** Cells give the F1 of each stage (one vs rest), the REM AUC, the F1 of N2+N3 and the N2+N3 AUC, for RF + HMM and SleepSEEG. N1 is poorly detected by every unit and classifier; thalamic N3 and REM F1 are low. Top (panel A), whole recording; bottom (panel B), night.

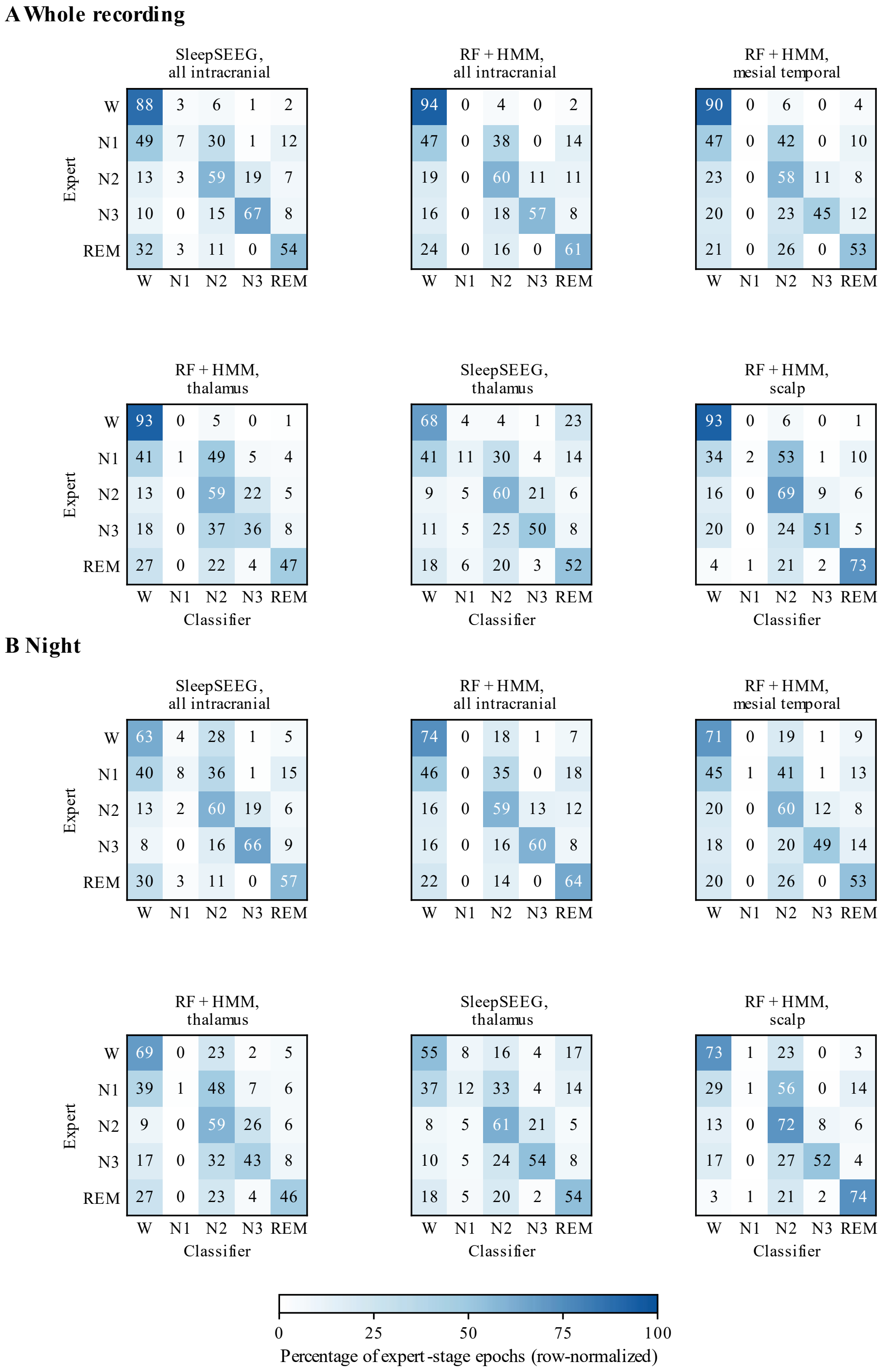


**Supplementary Figure S2. Pooled confusion matrices, normalized by expert stage (rows), for SleepSEEG on all intracranial channels and on the thalamus, and RF + HMM on all intracranial channels, the mesial temporal lobe, the thalamus and scalp.** Thalamic units confuse N2 and N3 and misclassify REM; SleepSEEG on the thalamus labels 23% of expert wake epochs as REM, vs 1% for RF + HMM (whole recording). Top (panel A), whole recording; bottom (panel B), night.

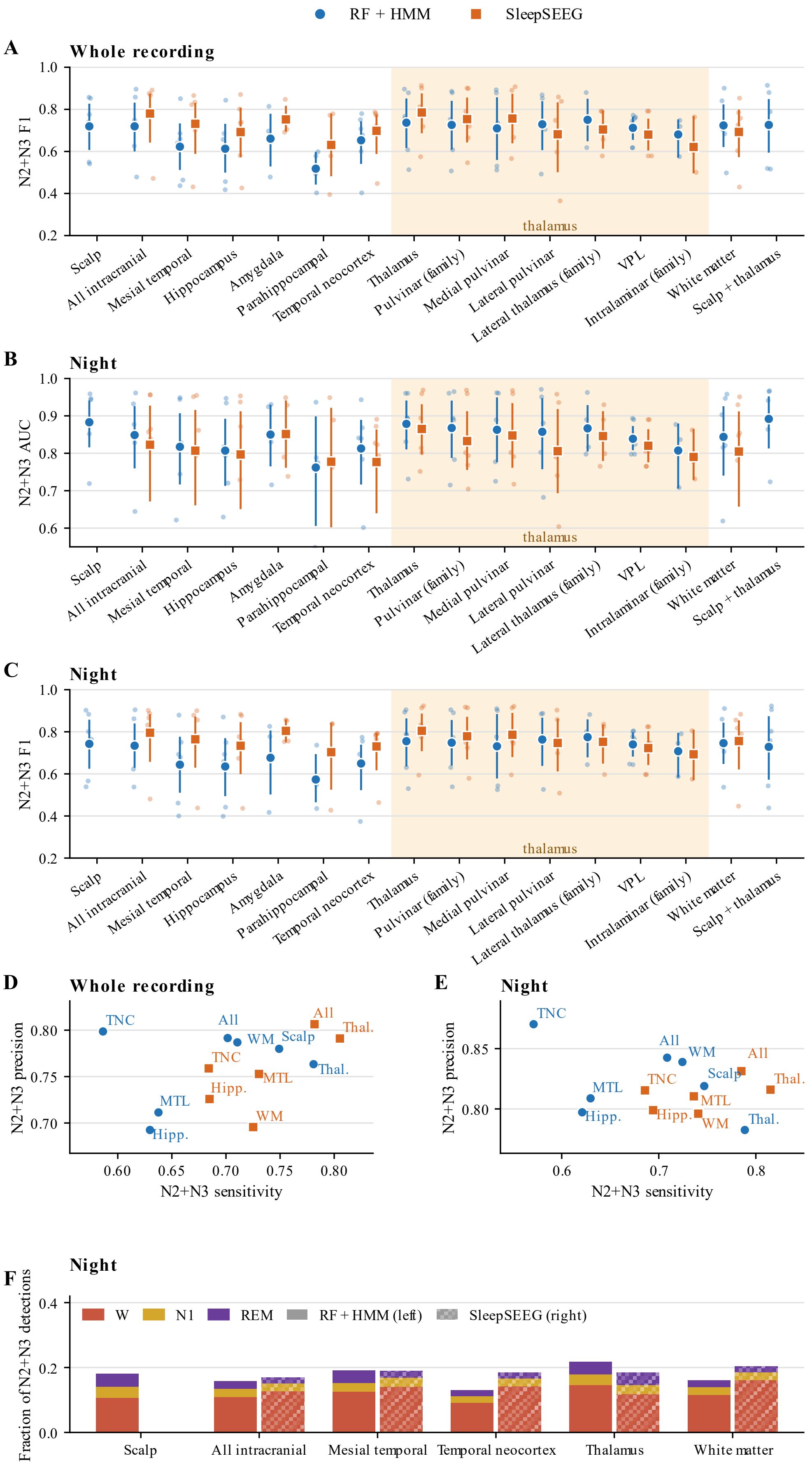

RF + HMM
SleepSEEG
A
Whole recording
N2+N3 F1
thalamus
Scalp
All intracranial
Mesial temporal
Hippocampus
Amygdala
Parahippocampal
Temporal neocortex
Thalamus
Pulvinar (family)
Medial pulvinar
Lateral pulvinar
Lateral thalamus (family)
VPL
Intralaminar (family)
White matter
Scalp + thalamus
B
Night
N2+N3 AUC
C
Night
N2+N3 F1
D
Whole recording
N2+N3 precision
N2+N3 sensitivity
TNC
All
WM
Scalp
Thal.
MTL
Hipp.
E
Night
F
Night
Fraction of N2+N3 detections
W
N1
REM
RF + HMM (left)
SleepSEEG (right)

**Supplementary Figure S3. Detection of N2+N3 sleep across regions: measures not shown in Figure 2**. **(A)** F1 of the decoded N2+N3 labels over the whole recording and **(B)** AUC for N2+N3 versus all other stages and **(C)** F1 at night, for scalp, the full intracranial montage, the mesial temporal lobe and its structures, the temporal neocortex, the thalamus and its nuclei, the white-matter control and scalp + thalamus fusion, for RF + HMM (circles) and SleepSEEG (squares). Small dots are participants; markers and bars are the mean and bootstrap 95% CI across participants. Shading marks thalamic units; SleepSEEG is not defined for scalp or fusion. **(D, E)** Sensitivity and precision of N2+N3 detection by unit (mean across participants) over the whole recording **(D)** and at night **(E)**. **(F)** Composition of false N2+N3 detections at night: fraction of the epochs each classifier labeled N2+N3 that the expert scored as W, N1 or REM (left bars, RF + HMM; right, hatched bars, SleepSEEG), mean across participants. All, all intracranial channels; Hipp., hippocampus; MTL, mesial temporal lobe; TNC, temporal neocortex; Thal., thalamus; WM, white matter.

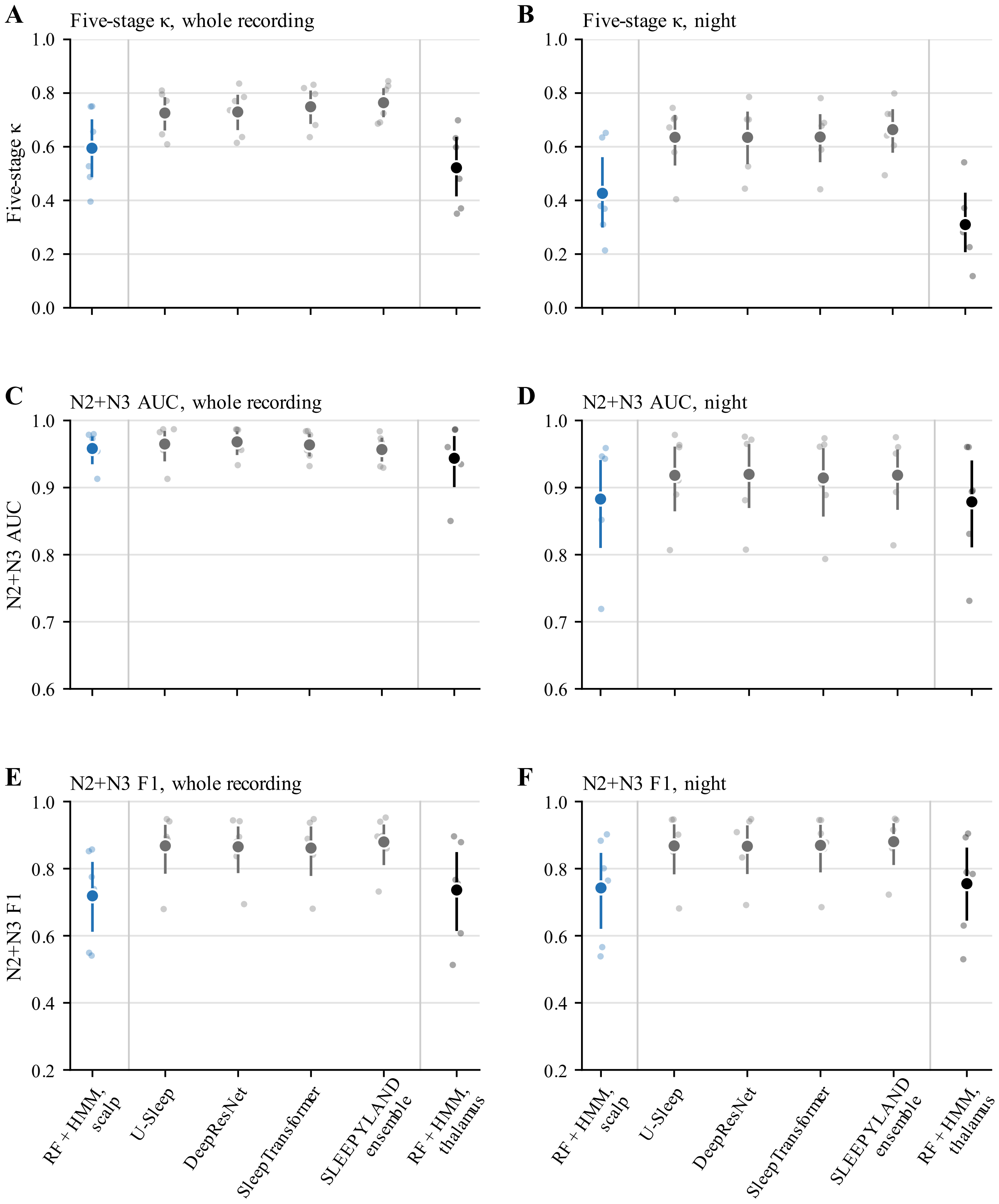


**Supplementary Figure S4. Pretrained scalp scorers. (A, B)** Five-stage κ, **(C, D)** N2+N3 AUC and **(E, F)** N2+N3 F1 against the expert for RF + HMM on scalp (blue; the scalp reference of the main analysis), the SLEEPYLAND scorers applied to the FCz-referenced scalp EEG (U-Sleep, DeepResNet, SleepTransformer**,** the SLEEPYLAND ensemble; gray) and, for reference, RF + HMM on the thalamus (black). Small dots are participants (recordings averaged within participant); markers and bars are the mean and bootstrap 95% CI across participants. Left (A, C, E), whole recording; right (B, D, F), night.

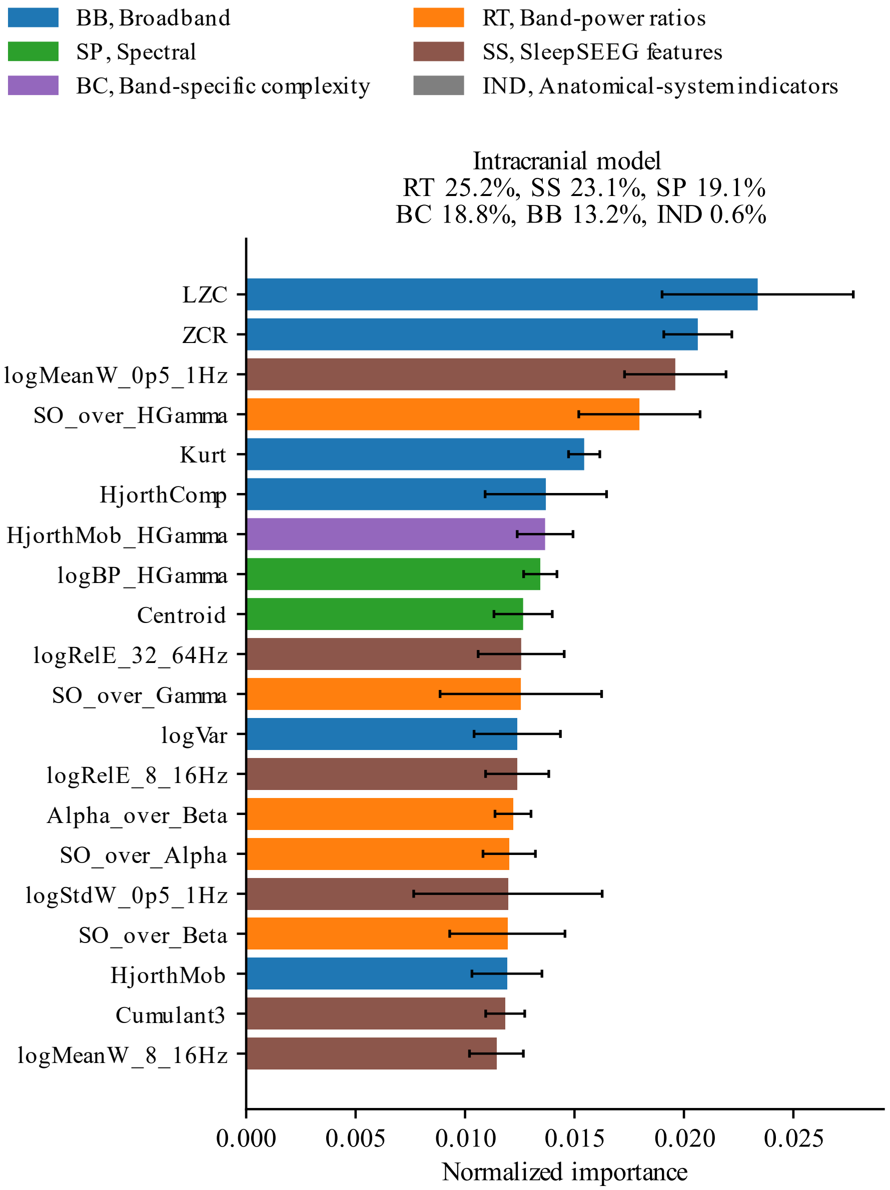


**Supplementary Figure S5. Random-forest predictor importance.** The 20 most important features of the intracranial model, normalized to the model's total importance and averaged across leave-one-participant-out folds (bars, mean; whiskers, SD). Colors and the percentages above the panel give feature families: BB, broadband; BC, band-specific complexity; IND, anatomical-system indicators; RT, band-power ratios; SP, spectral; SS, SleepSEEG features. LZC, Lempel–Ziv complexity; ZCR, zero-crossing rate. Importance reflects use by the trees.

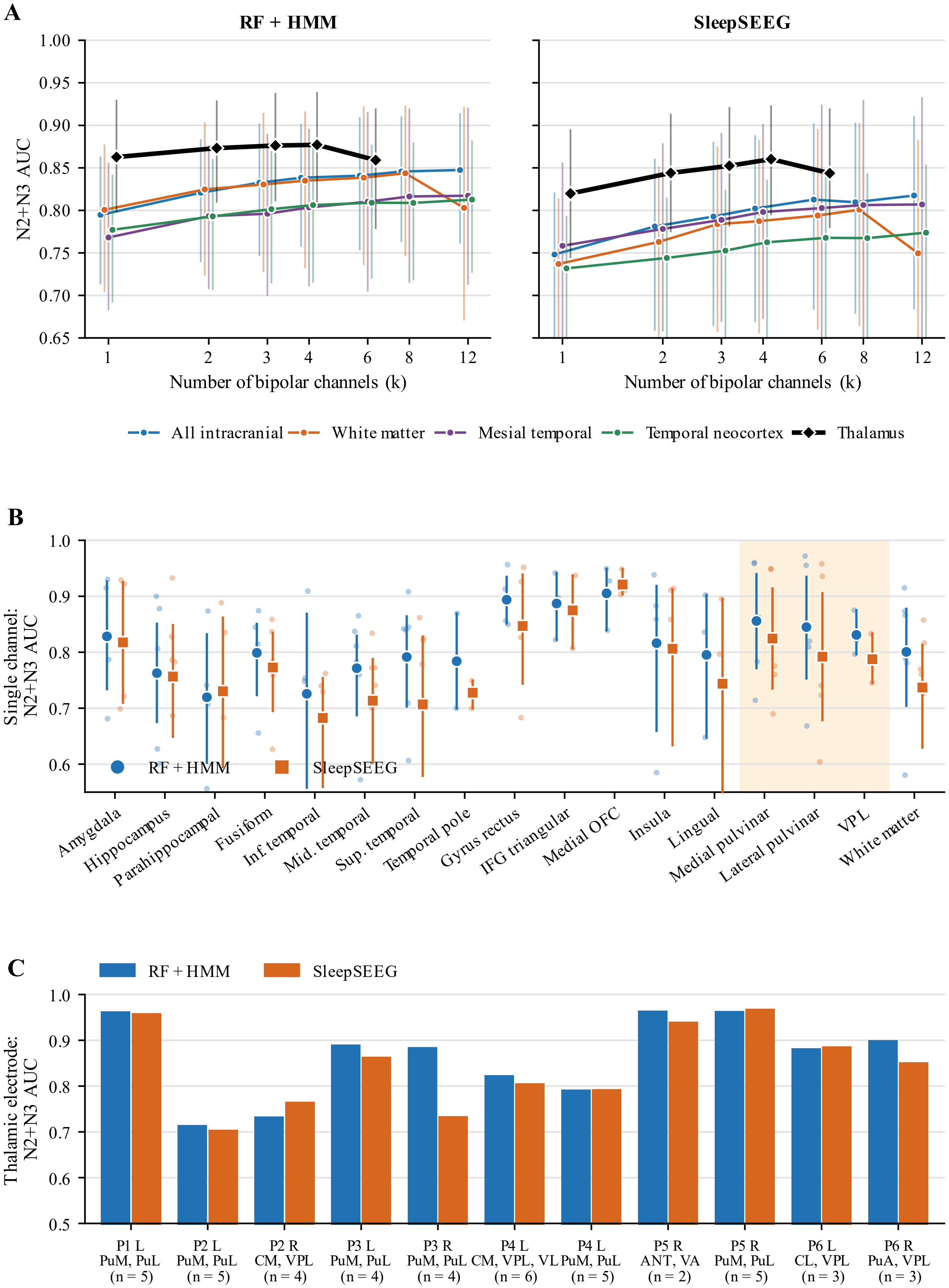
A
RF + HMM
SleepSEEG
N2+N3 AUC
1.00
0.95
0.90
0.85
0.80
0.75
0.70
0.65
1
2
3
4
6
8
12
Number of bipolar channels (k)
All intracranial
White matter
Mesial temporal
Temporal neocortex
Thalamus
B
Single channel: N2+N3 AUC
1.0
0.9
0.8
0.7
0.6
RF + HMM
SleepSEEG
Amygdala
Hippocampus
Parahippocampal
Fusiform
Inf. temporal
Mid. temporal
Sup. temporal
Temporal pole
Gyrus rectus
IFG triangular
Medial OFC
Insula
Lingual
Medial pulvinar
Lateral pulvinar
VPL
White matter
C
RF + HMM
SleepSEEG
Thalamic electrode: N2+N3 AUC
1.0
0.9
0.8
0.7
0.6
0.5
P1 L PuM, PuL (n = 5)
P2 L PuM, PuL (n = 5)
P2 R CM, VPL (n = 4)
P3 L PuM, PuL (n = 4)
P3 R PuM, PuL (n = 4)
P4 L CM, VPL, VL (n = 6)
P4 L PuM, PuL (n = 5)
P5 R ANT, VA (n = 2)
P5 R PuM, PuL (n = 5)
P6 L CL, VPL (n = 3)
P6 R PuA, VPL (n = 3)

**Supplementary Figure S6. Channel-count control and detection from single channels and single electrodes, night.** Panels and conventions as in Figure 3, for the night.
